# Impact of intragenic *NRXN1* deletions on early cortical development

**DOI:** 10.1101/2024.08.26.609494

**Authors:** Lucia Dutan, Nicholas J. F. Gatford, Thaise Carneiro, Roland Nagy, Victoria Higgs, APEX Consortia, Aicha Massrali, Arkoprovo Paul, Maria Fasolino, Zhaolan Zhou, Frances A Flinter, Dwaipayan Adhya, Maja Bucan, Simon Baron-Cohen, Deepak P. Srivastava

**Affiliations:** Department of Basic and Clinical Neuroscience, Maurice Wohl Clinical Neuroscience Institute, Institute of Psychiatry, Psychology and Neuroscience, King’s College London, London, UK, SE5 9NU, UK; MRC Centre for Neurodevelopmental Disorders, King’s College London, London, UK; Department of Genetics, Perelman School of Medicine at the University of Pennsylvania, Philadelphia, 19104-6145 USA; Autism Research Centre, Department of Psychiatry, University of Cambridge, Cambridge, CB2 8AH UK; Department of Clinical Genetics, Guy’s and St. Thomas’ NHS Foundation Trust, London, United Kingdom

**Keywords:** *NRXN1*isoforms, Intronic deletions, Synaptic function, Dendritogenesis, Neurodevelopmental disorders, Sex differences

## Abstract

**Background:** Deletions in *NRXN1* are strongly associated with neurodevelopmental and psychiatric conditions. While exonic deletions are well-studied, intragenic deletions, particularly in intron 5, are less understood and generally consider benign. Recent studies show exonic deletions impact isoform diversity during neurodevelopment, affecting neurogenesis and neuronal function. However, whether intragenic deletions impact isoform expression and neurodevelopment remains underexplored.

**Methods:** We used hiPSCs from typically developing individuals (control) and those with *NRXN1* intron 5 deletions to study neurodevelopment. HiPSCs were differentiated towards a cortical fate, with *NRXN1* isoform expression, molecular differences, and neuronal morphology examined.

**Results:** We observed distinct *NRXN1* isoform expression dynamics during early neurodevelopment, with two expression peaks post-neuronal induction and *NRXN1β* being most highly expressed. Both *NRXN1* deletion and control lines showed similar acquisition of regional and cell fate identity, but significant differences in *NRXN1* isoform expression were observed between deletion and control lines, and between deletion lines. RNA sequencing revealed genotype-dependent alterations, particularly in pathways related to synaptic function and neuronal morphology. Consistent with these findings, *NRXN1* deletion lines exhibited altered dendrite outgrowth, with variations between deletion lines.

**Conclusions:** Our results indicate a potential role for intron 5 in controlling *NRXN1* isoform expression during neurodevelopment. Alterations in gene expression profiles, correlated with morphological changes, suggest a role for *NRXN1* isoforms in shaping dendritic morphology. Molecular and cellular differences observed between lines with identical intronic deletions suggest that additional factors, such as genetic background or biological sex, may also play an important role in these phenotypes. Collectively, these findings indicate that *NRXN1* intronic deletions are not benign, influencing isoform expression, cellular phenotypes, and neurodevelopment.

## Introduction

Neurexin 1 (NRXN1) is a presynaptic cell adhesion molecule that interacts with postsynaptic proteins to regulate synapse organization and transmission (Pak et al., 2015; Shehhi et al., 2018; Hu et al., 2019, Boxer & Aoto, 2022). Both heterozygous and homozygous deletions in the *NRXN1* gene have been associated with various neurodevelopmental and psychiatric conditions, including autism spectrum conditions (herein referred to as autism), intellectual disability, and schizophrenia (Pak et al., 2015; Shehhi et al., 2018; Kasem et al., 2018; Molloy et al., 2023). Exonic deletions in *NRXN1,* particularly in exons near the 5ʹ end, have been associated with an increased likelihood of autism and schizophrenia (Tromp et al., 2021). In contrast, intronic deletions, although common, do not show the same level of association with clinical phenotypes (Lowther et al., 2017). Nevertheless, there are multiple reports where intragenic deletions in *NRXN1*, particularly within intron 5, have reported association with a range of neurodevelopmental conditions (Ching et al., 2010; Schaaf et al., 2012; Béna et al., 2013; Curran et al., 2013). Despite these reports, whether and how intragenic *NRXN1* deletions impact the cellular and molecular function of the gene remains underexplored (Kim et al., 2009; Lowther et al., 2017; Cooper et al., 2024).

The *NRXN1* gene spans 1.12 Mb, making it one of the largest genes in the human genome. The main isoforms of *NRXN1* are transcribed under the control of two independent promoters, generating a longer isoform, *NRXN1α*, and a shorter isoform, *NRXN1β* (Tabuchi & Südhof, 2002; Treutlein et al., 2014; Jenkins et al., 2015; Cooper et al., 2024) The *NRXN1α* promoter is located at the 5’ end of the gene, while the *NRXN1β* promoter is predicted to be located downstream of exon 17 (Wu el al., 2023; Xu et al., 2023; Cooper et al., 2024). Six canonical splicing sites have been described for *NRXN1α*, including sites 4 and 5, which are also present in *NRXN1β*, leading to numerous *NRXN1* structural variants due to extensive alternative splicing (Treutlein et al., 2014; Harkin et al., 2016; Hu et al., 2019; Gomez et al., 2021; Fernando et al., 2023). In humans, RNA sequencing data from post-mortem brain tissue have shown that *NRXN1* transcripts are present in the cortex before synapse formation, specifically at weeks 8-12 after conception (Jenkins et al., 2015; Harkin et al., 2017). *NRXN1α* and *NRXN1β* isoforms are expressed at gestational week 14 during cortical development, with both isoforms exhibiting a peak of expression from the second trimester to postnatal age 3 (Jenkins et al., 2015). The *NRXN1* isoform repertoire determines NRXN1’s affinity for its binding partners, with specific isoform combinations providing distinctive synaptic functionalities (Schreiner et al., 2014; Fuccillo et al., 2015; Nguyen et al., 2015; Südhof, 2017; Fernando et al., 2023; Molloy et al., 2023).

Studies using human induced pluripotent stem cells (hiPSCs) have provided further insight into the early expression profiles of the main *NRXN1* isoforms and their functions during neuronal differentiation (Flaherty et al., 2019; Lam et al., 2019; Sebastian et al., 2023). Using both long- and short-read sequencing techniques, Flaherty and colleagues demonstrated that hiPSC models accurately reflects *NRXN1* transcript diversity in derived neurons compared to post-mortem foetal tissue (Flaherty et al., 2019). This approach identified 123 *NRXN1α* isoforms in hiPSC-derived neurons, similar to that seen in human tissue. HiPSCs generated from individuals harbouring deletions in the gene have also begun to reveal differences in isoform expression and potential links with cellular and molecular phenotypes. For instance, differentiation of hiPSCs generated from an autistic individual with a homozygous deletion in *NRXN1α*, resulted in the generated a higher proportion of differentiated astroglia cells and immature neurons compared to controls, with neurons exhibiting reduced calcium signalling activity and impaired maturation of action potentials (Lam et al., 2019). Similarly, hiPSC-derived organoid models have also shown that exonic *NRXN1* heterozygous gene variants lead to alterations in synaptic signalling in maturing glutamatergic and GABAergic neurons, indicating that *NRXN1* is crucial for neuronal differentiation and the maturation of functional excitatory and inhibitory neurons (Sebastian et al., 2023). Patient cell lines with heterozygous exonic deletions at the 5’ or 3’ regions of *NRXN1* demonstrate changes in the diversity and transcript abundances of *NRXN1* isoforms within the derived neurons (Flaherty et al., 2019; Fernando et al., 2023). These neurons exhibited a reduction in non-deleted isoforms and the generation of novel isoforms, which could be mimicked or attenuated by altering the expression of wildtype or novel isoforms, demonstrating a relationship between *NRXN1* isoforms and cellular phenotypes (Flaherty et al., 2019; Fernando et al., 2023). Recent work from our group demonstrated that patient lines with heterozygous deletions located either within intron 5, or including intron 5 and surrounding exons, produced similar alterations in cortical neurogenesis (Adhya et al., 2021). Both cell lines, generated from individuals diagnosis with autism or developmental delay, demonstrated altered cell fate acquisition in addition to impaired neural rosette formation, cellular phenotypes similarly observed in patient lines generated from idiopathic autistic individuals. Interestingly, unsupervised clustering approaches separated patient lines based on cellular phenotypes; with both hiPSC lines with overlapping deletions in intron 5, clustering together (Adhya et al., 2021). These findings indicate that deletions in intron 5 of *NRXN1* may uniquely contribute to cellular phenotypes relevant for neurodevelopmental conditions, but how this occurs at the cellular and molecular level is unclear.

In this study, we aimed to investigate the impact of intragenic deletions within *NRXN1* on cellular and molecular phenotypes in developing cortical neurons. Using hiPSCs generated from related individuals carrying identical deletions in intron 5 as well as from an unrelated individual with an overlapping but larger deletion, we observe that intragenic deletions alter the expression of major *NRXN1* isoforms throughout cortical differentiation. In addition, we find that intragenic *NRXN1* deletions change the gene expression profile of immature cortical neurons, as well as altering the overall dendritic morphology of these cells. Together, these data indicate that *NRXN1* intragenic deletions play a complex role in regulating isoform expression, which in turn impacts the development of cortical neurons.

## Materials and Methods

### Human induced pluripotent stem cells (hiPSC) culture and neuronal differentiation

Human induced pluripotent stem cells (hiPSCs) were participants recruited from the Longitudinal European Autism Project (LEAP) (Loth et al., 2017) or Brain and Body Genetic Recourse Exchange (BBGRE) studies (Ahn et al., 2013) via the StemBANCC Innovative Medicines Initiative or the EU-AIMS Innovative Medicines Initiative. Recruitment of participants was conducted in accordance with the guidelines of the ‘Patient hiPSCs for Neurodevelopmental Disorders (PiNDs) study’ (REC No 13/LO/1218). Autism diagnosis was determined based on Autism Diagnostic Observation Schedule or Autism Diagnostic Interview–Revised scores, while neurotypical control subjects were selected from the population if they had no diagnosis of any psychiatric condition. Diagnostic details can be found in our previous study (Adhya et al., 2021). Reprogramming was performed as previously described (Cocks et al., 2014; Adhya et al., 2021) using the CytoTune-iPS 2.0 Sendai Reprogramming Kit (Thermo Fisher, A16517) with the transcription factors KLF4, SOX2, C-MYC, and OCT4. The derived cell lines included hiPSCs from unrelated neurotypical (control) participants – female 007_CTF and males CTR_M3, 127_CTM – and proband cell lines from a neurotypical female, 110_NXF and her son 109_NXF who had a diagnosis of autism and microcephaly, both carrying an identical 60Kb heterozygous deletion in *NRXN1* intron 5. Cell lines from an additional female, 092_NFX, with a diagnosis of developmental delay and harbouring a 200Kb heterozygous deletion spanning from intron 5 to exon 8 of the *NRXN1* gene, was also used in this study (**Figure 1A**). Experiments were conducted using three independent clones per cell line, except for control lines 007_CTF and 127_CTM, where only one clone was used.

**Figure 1:**
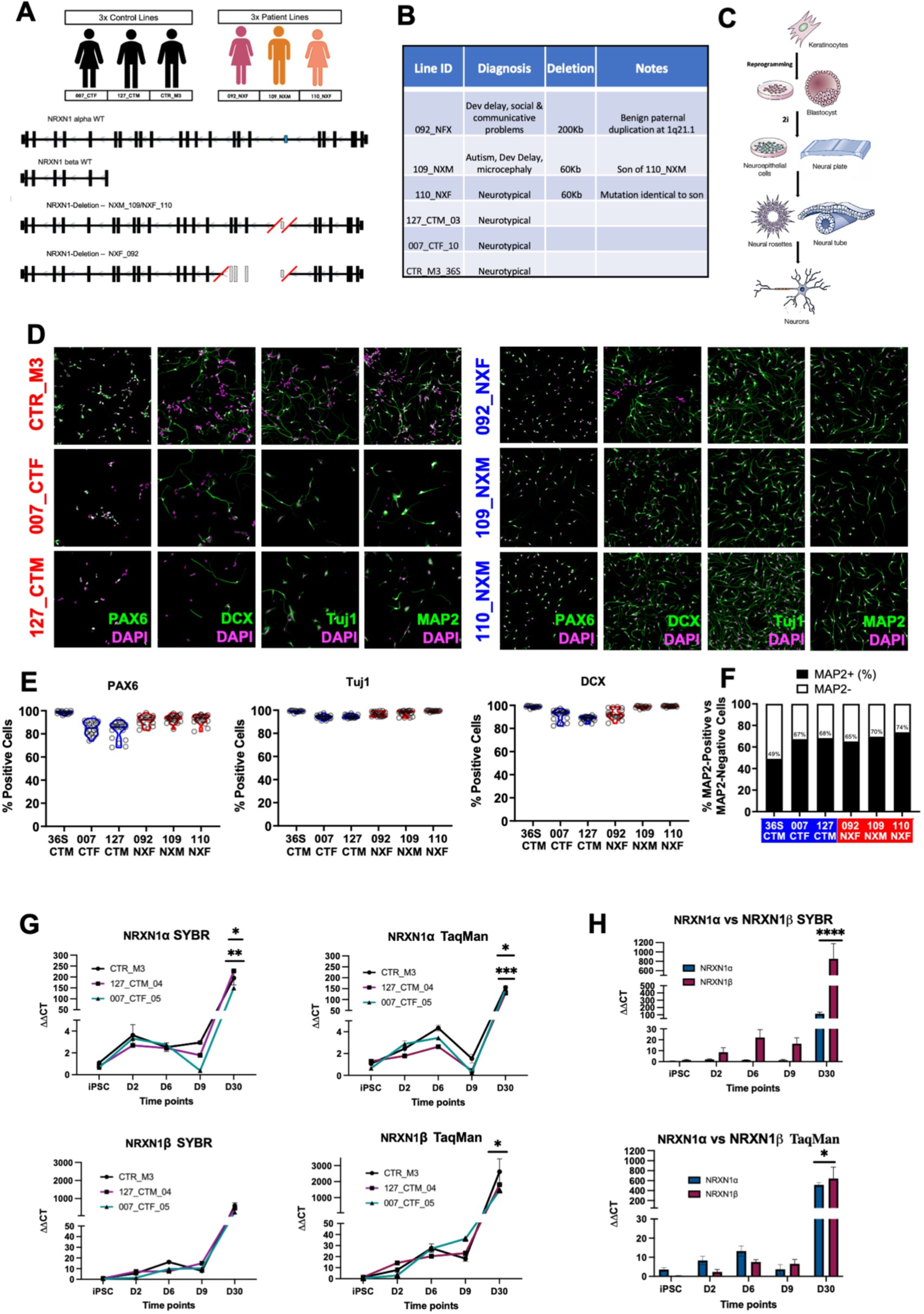
Characterization of Patient Cohort and *NRXN1* Deletions hiPSC lines. **(A)** Schematic representation of the study cohort, comprising three neurotypical controls and three individuals with *NRXN1* deletions. Genetic deletions in *NRXN1* are depicted with red boxes, with coordinates based on the University of California Santa Cruz (UCSC) Genome Browser. **(B)** Detailed overview of *NRXN1* deletions in the patient cohort, showing participant diagnoses and deletion sizes in kilobases (Kb). **(C)** Schematic of the neuronal differentiation process using 2i, mapping stages from hiPSC to various stages of embryonic development: from pluripotent cells resembling epiblast cells, through neuroepithelial and neural rosette stages, to immature neurons. **(D-E)** Immunocytochemical analysis of neuronal marker expression (PAX6, DCX, TUBB3 (Tuj1), and MAP2) in control and *NRXN1* deletion lines at day 23 post-differentiation, with quantification showing the percentage of cells expressing each marker. Geometric means are: PAX6 (88.79±4.88%), DCX (92.84±3.06%), TUBB3 (Tuj1) (95.94±1.61%), and MAP2 (61.52±6.24%) in controls; PAX6 (92.26±0.46%), DCX (96.68±2.19%), TUBB3 (Tuj1) (97.85±0.78%), and MAP2 (69.44±2.45%) in *NRXN1* deletion lines. **(F)** Ratiometric comparison of MAP2-positive versus MAP2-negative cells between control (blue) and *NRXN1* deletion (red) lines, indicating no significant differences. **(G)** Temporal expression patterns of *NRXN1α* and *NRXN1β* in control lines (CTR_M3, 127_CTM and 007_CTF) over developmental time points (hiPSC, d2, d6, d9, and d30), analysed using SYBR and TaqMan assays. The expression of *NRXN1α* and *NRXN1β* isoforms in the cell lines exhibits peaks on days 6 and 30. On day 30, significant differences are evident in the expression of *NRXN1α* SYBR between CTR_M3_36S and 007_CTF, as well as between 127_CTM and 007_CTF. Similarly, *NRXN1α* TaqMan analysis indicates significant differences between CTR_M3_36S and 127_CTM, and between 127_CTM and 007_CTF at the same time point. For *NRXN1β* TaqMan, significant differences are observed at day 30 between CTR_M3_36S and 007_CTF_05. **(H)** comparative analysis of *NRXN1α* and *NRXN1β* across five developmental stages shows a consistently higher expression profile for *NRXN1β*. At day 30, significant differential expression is detected between *NRXN1α* and *NRXN1β* using both SYBR and TaqMan assays, reinforcing the trend of higher NRXN1β activity throughout the developmental stages.

The cells were cultured in StemFlex media (Thermo Fisher, A3349401) in Nunc treated 6-well multidishes coated with 2% Geltrex (Life Technologies, A1413302), with media replaced daily. Upon reaching confluence, cells were differentiated using N2:B27 neural induction medium composed of N2 supplement (Life Technologies, 17502-048) at 1X final concentration in DMEM (Sigma, D6421) with 1X Glutamax (Gibco, 35050061) + B27 supplement (Life Technologies, 17504-044) at 1X final concentration in Neurobasal Media (Life Technologies; 21103-049) with 1X Glutamax. Small molecule inhibitors SB431542 (10µM, Sigma, S4317) and LDN193189 (0.1µM, Sigma, SML0559) were added to facilitate induction (Shum et al., 2020). After 8 days, the inhibitors were discontinued, and cells were maintained in N2:B27 media, passaged at days 8, 12, 15, and 18 using StemPro Accutase (A1110501), and replated at a 1:1 ratio in media containing Rho-kinase inhibitor (10 µM, Sigma, SCM075). Terminal differentiation was initiated on day 20, with cells seeded in T25 flasks or 96-well plates for RNA extraction and immunocytochemistry, respectively, coated with poly-D-lysine 5 µg/ml (Sigma, P6407) and laminin 20 µg/ml (Sigma, L2020). After day 20, the media was replaced with B27 media supplemented with 200 μM L-ascorbic acid (Sigma, A4544) and 10 μM DAPT (Santa Cruz Biotechnology, sc-201315A). DAPT was discontinued after day 27. RNA was harvested at time points day 0 (hiPSC) and days 2, 6, 9 and 30 after neural the induction. Neurons were differentiated for 24 days before fixing for immunocytochemistry and high content screening assays.

### RNA extraction and Complementary DNA synthesis

Total RNA was harvested and lysed with Trizol reagent (Life technologies, 15596026) and isolated by centrifugation with 200 μl of 100% Chloroform followed by 500 μl of 100% isopropanol and 1ml of 80% ethanol. The RNA pellet was diluted with 30μl of water and incubated with 3 volumes of Ethanol 80% and 0.1 volumes of Na-Acetate at −80°C overnight. The RNA was pelleted by centrifugation and cleaned with 2 washes with 1ml of Ethanol 80%. The RNA was purified by air drying the pellet and diluting it with 30μl of water. The quality control and concentration were assessed by analyzing the RNA with the NanoDrop 1000 Spectrophotometer (Thermo scientific). Complementary DNA (cDNA) was synthetized from 1μg of RNA using SuperScript III Reverse Transcriptase (Thermo Fisher, 18080093) following manufacturer’s instructions with oligo(dT) (Thermo Fisher, 18418020) priming.

### Quantitative Real-Time PCR

A set of 12 primers were designed using gene sequences extracted from the University of California and Santa Cruz (UCSC) genome and bioinformatics browser Primer3Plus (v. 0.4.0) software (**Table 1**). The primers designed for *NRXN1α* transcripts target exon 7 and exon 8, while the primers for *NRXN1β* are designed to amplify the region spanning intron 17 of *NRXN1α*, which constitutes exon 1 for *NRXN1β*, and exon 18 of *NRXN1α* **(Supplementary Figure 1)**. The real-time PCR analyses were conducted by using 2μl of cDNA per well in 384 well plates (Applied Biosystems, 4309849) and adding 0.3μM primers mix and HOT FIREPol EvaGreen qPCR Mix Plus (ROX) (Solis biodyne, 08-24-00020) to a 1X final concentration. The reaction was run in a QuantStudio 7 Real-Time PCR thermocycler detection system.

PCR reaction conditions: 95°C for 12m for the initial activation followed by 40 cycles of 95°C for 15s, 60°C for 20s and 72°C for 20s. The melting curve analyses were preformed from 60°C to 95°C with readings every 1°C. The housekeeping genes *GAPDH*, *SDHA* and *RPL27* were used to normalize the genes expression. Additionally, the gene expression levels of *NRXNα*, *NRXN1β*, and *pan NRXN1* were quantified using specific TaqMan probes labelled with FAM dye (Applied Biosystems, Hs00985129_m1, Hs00373346_m1 and Hs00985123_m1 respectively), using the housekeeping gene *GAPDH* (Applied Biosystems, 4310884E) as endogenous control (VIC). All assays were performed according to the manufacturer’s protocols (Applied Biosystems) using 1ul of each TaqMan Gene Expression Assay and 10 μl TaqMan Fast Advanced Master Mix (Applied Biosystems, 444556).

The ΔΔCt comparative method of relative quantification was used to quantify gene expression at different time points using an average of the time point hiPSC controls as a reference sample for all reactions. Each sample was run in triplicate and the 3 CT values per clone were averaged and used to generate the relative gene expression values per cell line. The means between samples were compared by a two-way ANOVA test with Tukey’s correction with 95% confidence interval (CI) from time points iPSC to day 9 and with a one-way ANOVA Tukey’s correction with 95% CI at day 30 with Prism package of GraphPad software.

**Table1.**
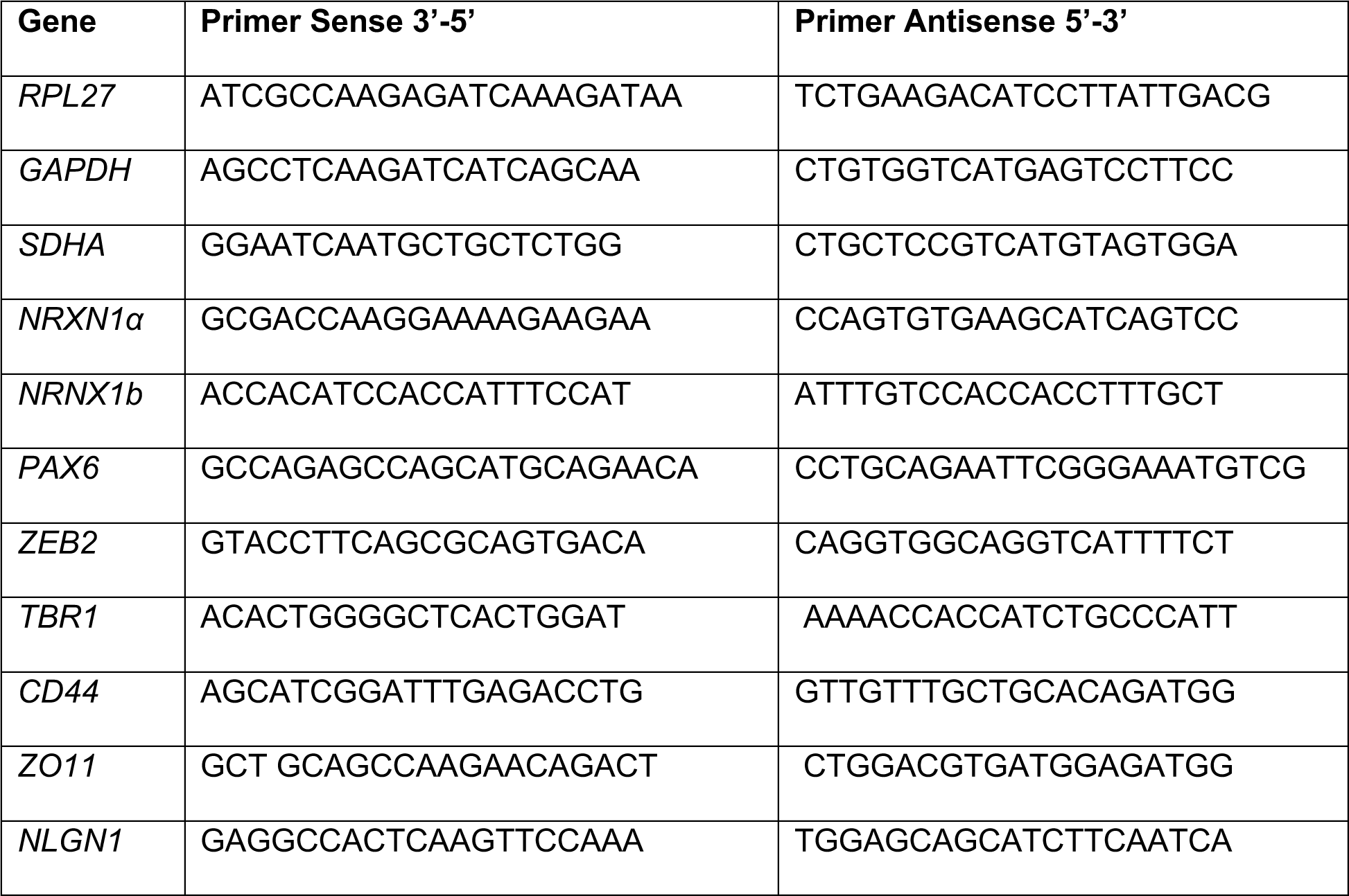
List of primer pairs used for Q-PCR.

### Immunocytochemistry

Cells were fixed at time points hiPSC, day 4 and day 8 and day 24 for immunocytochemistry assays with 4% formaldehyde/4% sucrose in PBS for 20 minutes at RT. Cells were subsequently rinsed twice in PBS, then permeabilized and blocked simultaneously in PBS containing 0.1% Triton-X-100 and 2% Normal Goat Serum (NGS) (Cell Signaling Technology, 5425S) for 1 hour at RT. Primary antibodies were added to each well in PBS containing 2% NGS and incubated overnight at 4°C (**Table 2**). Wells were washed 3 times with PBS for 15 minutes each time. Alexa Fluor 488/568/633 conjugated secondary antibodies in PBS containing 2% NGS were added to each well for 1 hour at RT, followed by two PBS rinses and three 15-minute PBS washes at RT. Once washed, all wells were incubated for 5 minutes in PBS containing 0.1% 4′,6-diamidino-2-phenylindole (DAPI) (Life Technologies, D1306) to counterstain nuclei. Image acquisition was performed using Cell insight CX5 Hight Content Screen Platform (Thermo Fisher, CX51110) with a 10X (day 0 and day 24) or 20X (day 4 and day 8) objective. The expression of specific proteins was quantified by using the bioapplication Cell Health Profiling from the iDev software package (Thermo Fisher). Nuclear staining was used to identify valid cells. Specific parameters of shape, cell location, size and the intensity were used to identify positive and negative cells. Non-specific background was determined by using purified mouse IgG (Merck Millipore, cs200621) and purified rabbit IgG instead of primary antibodies. The positive cells percentage at each time point were compared by a one-way ANOVA.

**Table 2.** List of antibodies used for immunocytochemistry.

| **Antibody** | **Type** | **Host** | **Supplier** | **Catalog Number** | **Clone Number** |
| --- | --- | --- | --- | --- | --- |
| TUJ1 | Monoclonal | Mouse | Biolegend | 801202 | TUJ1 |
| AFP | Monoclonal | Mouse | Invitrogen | MA5-14666 | P5B8 |
| SMA | Polyclonal | Rabbit | Abcam | ab5694 | N/A |
| Nanog | Polyclonal | Rabbit | Abcam | ab80892 | N/A |
| OCT4 | Monoclonal | Rabbit | Invitrogen | 701756 | 3H8L6 |
| SSEA4 | Monoclonal | Mouse | Invitrogen | MA1-021 | MC-813-70 |
| TRA-1-81 | Monoclonal | Mouse | Invitrogen | MA1-024 | TRA-1-81 |
| PAX6 | Polyclonal | Rabbit | BioLegend | 901301 | N/A |
| Nestin | Monoclonal | Mouse | BioLegend | 656802 | 10C2 |
| DCX | Polyclonal | Rabbit | Abcam | ab18723 | N/A |
| MAP2 | Polyclonal | Chicken | Abcam | ab92434 | N/A |

### RNA sequencing analysis

Bulk RNA sequencing was performed on 12 samples from three NRXN1 deletion lines (109_NXM, 110_NXF, and 092_NXF), each represented by three clones, and three control cell lines (127_CTM_03, 007_CTF_10, and CTR_M3_36S), each represented by one clone. This work was carried out at Genewiz in Azenta Life Sciences. Libraries were constructed using the NEBNext Ultra II RNA Library Prep Kit for Illumina (NEB), adopting a polyA selection strategy. RNA concentration was assessed using the Qubit 4.0 Fluorometer (Life Technologies), and RNA integrity was assessed with the Agilent 5300 Fragment Analyzer (Agilent Technologies). The libraries sequencing was completed on an Illumina NovaSeq 6000 using a 2×150 paired-end configuration. Base calling was conducted by the NovaSeq Control Software v1.7 and raw sequence data generated was converted into FastQ files using the Illumina bcl2fastq program.

Quality control of FastQ files was conducted using the FastQ Screen tool (Wingett & Andrews, 2018). Sequencing libraries were aligned to the human reference genome (GRCh38.p13) using the STAR aligner (Dobin et al., 2013) with default settings. The aligned sequences were then sorted by chromosomal coordinates using the samtools sort function (Danecek et al., 2021). Duplicate reads were identified and tagged using Picard’s MarkDuplicates tool. Gene counts were quantified using the HTseq-count function from the HTseq package (Anders et al., 2015), which facilitated the identification of reads overlapping with exonic regions of each gene. Differential expression analysis was conducted using the DESeq2 package in R/Bioconductor, applying default parameters to facilitate comparisons between two distinct groups. Specifically, analyses were performed between control cell lines and NRXN1 deletion lines, with additional targeted analyses conducted across different deletion regions. Significant differentially expressed genes (DEGs) were identified based on stringent criteria, requiring an adjusted p-value of less than 0.05 and a fold change exceeding 2. This approach allowed for the robust identification of genes significantly impacted by NRXN1 deletions.

Enrichment analyses of biological terms for the differentially expressed genes (DEGs) were conducted using the DAVID Gene Functional Classification Tool (Sherman et al., 2022). All DEGs served as the background dataset for these analyses, irrespective of their p-values or fold changes. The SynGO Ontologies platform was employed to investigate enriched synaptic terms within the sets of DEGs. This approach allowed for a detailed exploration of synaptic functions potentially impacted by the gene expression changes observed in the study. Additionally, overlaps between the identified DEGs and genes listed in SFARI gene categories 1 and 2 were examined using Fisher’s exact tests through the GeneOverlap package, with a 5% false discovery rate (FDR) correction applied to adjust for multiple comparisons. This methodological approach facilitated the visualization and statistical testing of gene overlaps, providing insights into the functional implications of the DEGs.

### Variant annotation and filtering workflow for Whole Genome Sequencing

We performed Whole Genome Sequencing (WGS) on three *NRXN1* deletion cell lines (092_NXF, 109_NXM, 110_NXF) and two control cell lines (007_CTF, 127_CTM) using the Illumina HiSeq platform with 2×150 PE and 30x coverage. Variants with a “PASS” filter, based on Variant Quality Score Recalibration (VQSR), and those with depth (DP) < 10 or genotype quality (GQ) < 20 were retained. We aligned sequences to the GRCh38.86 reference genome and annotated variants with ANNOVAR (Wang et al., 2010). Variants classified as intergenic, ncRNA_exonic, intronic, UTR3/UTR5, upstream/downstream, or synonymous SNVs were excluded. Non-synonymous SNVs, splicing variants, and gain/loss-of-function variants were selected for further analysis.

To prioritize variants with low to moderate minor allele frequencies, we excluded single nucleotide variants (SNVs) with a frequency above 0.1% using gnomAD v4.0 data (Karczewski et al., 2020). Pathogenicity was assessed using two tools: Rare Exome Variant Ensemble Learner (REVEL), which combines predictions from 13 algorithms and scores from 0 to 1, with higher scores indicating greater pathogenicity (loannidis et al., 2016), and Combined Annotation Dependent Depletion (CADD), which ranks SNVs and indels relative to all human genome variants. Variants with a CADD score of 10 or more fall within the top 10% of deleterious variants, while those with a score of 20 or above are in the top 1% (Kircher et al., 2014). Variants were selected for further analysis if they had a REVEL score ≥0.5 and/or a CADD score ≥20.

### Analytical approach for prioritizing and enriching rare genomic SNVs

We initiated our analysis by focusing on moderate and rare genetic variants, i.e. variants with minor allele frequency (MAF) of 0.1% or less, a CADD score of 20 or higher, and/or a REVEL score of 0.5 or greater. We compared a burden for these predicted deleterious variants, in the deletion cell lines (092_NXF, 109_NXM, 110_NXF) and controls (007_CTF, 127_CTM) cell lines. We employed a Welch two-sample t-test, which is designed to determine if there is a significant difference between the means of two independent groups while not assuming equal variances. The null hypothesis was tested for deviations in both directions to assess the significance of the observed differences.

After the initial burden analysis, we conducted a comprehensive assessment of these variants to evaluate their enrichment in specific pathways and gene ontologies related to biological processes. For this purpose, we utilized the DAVID database (https://david.ncifcrf.gov/) to analyse enrichment across various biological pathways and processes. This combined approach provided a thorough understanding of how the variants are associated with biological functions and synaptic processes.

Following our initial broad analysis, we narrowed our focus to examine two specific sets of autism high confidence genes: a) genes classified as level one and two by the Simons Foundation Autism Research Initiative (SFARI), which are recognized for their strong and moderate evidence of association with autism (https://genomes.sfari.org/); and b) autism high-risk genes identified through the Transmission and De Novo Association (TADA) test (Sanders et al. 2015; Feliciano et al. 2019; Ruzzo et al. 2019; Satterstrom et al. 2020; Fu et al. 2022), which provides a comprehensive evaluation of genetic risk factors based on transmission patterns and de novo mutations. This targeted approach allowed us to refine our analysis and gain more precise insights into the genetic underpinnings of autism by focusing on genes with robust evidence of involvement.

## Results

### Characterization of control and patient-hiPSC lines

HiPSCs cells were generated from hair keratinocytes as previously described (Adhya et al., 2021). Three hiPSC lines from a female (007_CTF) and 2 male (CTR_M3 and 127_CTM) typically developing individuals with no prior diagnosis of neuropsychiatric disorders were used in this study – these lines have previously been characterised (Adhya et al., 2021). Three patient lines from two females 092_NXF and 110_NXF and one male 109_NXM were also used in the study. HiPSC lines 109_NXM and 110_NXF are a mother/son pairing whereas 092_NXF was generated from an unrelated individual. All patients were recruited from the Brain and Body Genetic Recourse Exchange (BBGRE) study (Ahn et al., 2013). Microarray based comparative genomic hybridisation (CGH) revealed all three patient lines had overlapping deletion type copy number variant in the 2p16.3 region, specifically located with the *NRXN1* gene. HiPSC line 092_NXF carries a large heterozygous 200 kilobase (Kb) deletion in *NRXN1* starting at intron 5 and including exons 6, 7, and 8. Patient-hiPSC lines 109_NXM and 110_NXF carry smaller but identical heterozygous intronic 60kb deletions in *NRXN1* intron 5. The deletions in 109_NXM and 110_NXF overlapped entirely within the 200kb deletion found in 092_NXF (**Figure 1A**). The donor of cell line 109_NXM was diagnosed with autism, developmental delay, and microcephaly. In contrast, the donor of cell line 110_NXF, despite harbouring the same *NRXN1* deletion, had no diagnosis. The donor of cell line 092_NXF was diagnosed with developmental delay and social and communicative challenges (**Figure 1B**).

All hiPSC lines were assessed for pluripotency by examining the expression of pluripotency markers, alkaline phosphatase expression and the spontaneous formation of all three germ layers of the developing embryo by embryoid formation (Shum et al., 2020). Immunocytochemistry assays of control (M3_36S, 127_CTM and 007_CTF) and the three *NRXN1* deletion (110_NXF, 110_NXF and 092_NXF) lines were performed to demonstrate pluripotency **(Supplementary Figure 2)**. All lines show positive and similar levels of immunostaining for the pluripotency markers Nanog, OCT4, SSEA4, and TRA181; while showing no staining for neuronal progenitor markers NESTIN and PAX6 **(Supplementary Figure 2)**. The differentiation potential of each hiPSC line was validated through their spontaneous differentiation into cells representative of the three germ layers. This was achieved following the generation of embryoid bodies, as outlined in Cocks et al. (2014) **(Supplementary Figure 3).** These experimental approaches effectively confirmed the pluripotency and comprehensive differentiation capacity of the hiPSCs utilized in this study.

HiPSC lines were induced to acquire neural fate by mimicking embryonic developmental cues and inhibiting TGF and BMP with dual SMAD inhibition (2i) to induce the hiPSC to differentiate into neuroepithelial cells like the cells of the embryonic neural plate (Gupta et al., 2023). After 8 days of induction hiPSCs formed neural rosettes, that are equivalent to the cells of the neural tube; after 20 days of differentiation, cells are terminally plated to form immature neurons (**Figure 1C**). To confirm the neuronal differentiation potential of all hiPSCs, cell lines were differentiated until day 23 and assessed for the expression of pan neuronal markers PAX6, DCX, TUBB3 (Tuj1), MAP2 using a high content screening platform. Immunocytochemistry analysis of the hiPSC-derived neurons from control and *NRXN1* deletion lines demonstrated that all lines were positive and had similar levels of expression of PAX6, DCX, TUBB3 (Tuj1), MAP2 (**Figure 1D-E**). Ratiometric analysis comparing MAP2-positive vs MAP2-negative cells per line revealed the majority of cell lines produced ∼70% MAP2-positive cells at this early stage of cortical differentiation (**Figure 1F**) with no significant differences observed between cell lines. These data suggest that control and *NRXN1* deletion lines efficiently differentiated into cortical neurons in a similar manner.

### *NRXN1* isoform temporal expression profile across cortical differentiation

Previous studies have established that *NRXN1* and its principal isoforms are expressed in foetal brain tissue (Jenkins et al., 2015) and in hiPSC-derived cells differentiating towards a cortical fate (Sebastian et al., 2023). To further explore the expression dynamics of *NRXN1* isoforms during early neuronal differentiation, three control hiPSC lines were directed towards a forebrain cortical fate. The expression of the main isoforms, *NRXN1α* and *NRXN1β*, was monitored over 30 days of neural induction at specific time points (hiPSC, days 2, 6, 9, and 30) using quantitative real-time PCR (Q-PCR). For detection, we employed specific primers and TaqMan probes (Jenkins et al., 2015) targeting specific regions of the *NRXN1* isoforms (**Supplementary Figure 1**). A significant increase in *NRXN1* expression was observed 2 days post-induction, with transcript levels peaking at days 6 and 30 (**Figure 1G**). These observations suggest a potential role for *NRXN1* in early neuronal development. The expression profiles of *NRXN1α* and *NRXN1β* were consistent across all control cell lines during the initial 9 days of differentiation, although the use of different detection methods (SYBR versus TaqMan) introduced slight variations in the measured expression levels (**Figure 1G**). These differences are likely due to the diverse *NRXN1* isoform repertoire targeted by the primers designed for SYBR or TaqMan amplification **(Supplementary Figure 1)**.

In immature neurons (day 30), significant differences in the expression of *NRXN1α* SYBR between CTR_M3_36S and 007_CTF (Two-way ANOVA: *padj = 0.0456, 95% CI) and between 127_CTM and 007_CTF (Two-way ANOVA: **padj = 0.0073) were observed. Similarly, *NRXN1α* TaqMan analysis shows significant variability at day 30 between the cell lines CTR_M3_36S and 127_CTM (Two-way ANOVA: ***padj = 0.0008, 95% CI) and between 127_CTM and 007_CTF (Two-way ANOVA: *padj = 0.0202). For *NRXN1β* TaqMan, significant differences are noted at day 30 between the lines CTR_M3_36S and 007_CTF_05 (Two-way ANOVA: **padj = 0.0457, 95% CI). Further analysis revealed differential expression profiles between the isoforms. *NRXN1β* isoforms were consistently upregulated compared to *NRXN1α* across all stages of neuronal differentiation, mirroring findings from post-mortem analyses (Jenkins et al., 2015). At day 30, significant differential expression between *NRXN1α* and *NRXN1β* was observed using SYBR (Two-way ANOVA: ****padj < 0.0001, 95% CI) and TaqMan assays (Two-way ANOVA: *padj = 0.0107, 95% CI) (**Figure 1H**).

### Impact of heterozygous intronic deletions on *NRXN1* isoforms across cortical differentiation

*NRXN1*, known for its role as an adhesion molecule in synapse formation and maintenance, has also been identified to be active during early stages of cortical differentiation (Lam et al., 2019; Sebastian et al., 2023). Consistent with these findings, we have previously shown that early corticogenesis and formation of neural rosettes are altered in cell lines containing heterozygous intron 5 deletions (Adhya et al., 2021). Therefore, we carried out a deeper investigation into *NRXN1* isoform expression across various stages of cortical development in both control lines and *NRXN1* deletion lines. We first confirmed that hiPSCs carrying deletions in intron 5 of *NRXN1* displayed altered rosette formation. Similar to what we have previously reported (Adhya et al., 2021), differentiation of cell line 109_NXM resulted in smaller neural rosettes at day 8 compared to those generated by control lines (**Supplementary Figure 4**). Next, we assessed the expression patterns of *NRXN1* isoforms *panNRXN1*, *NRXN1α*, and *NRXN1β* expression in control and *NRXN1* deletion lines at key developmental time points (hiPSC, day 2, day 6, and day 9 - **Figure 2A & B**) using *NRXN1* isoform specific primers (SYBR - **Supplementary Figure 1**). Expression analysis revealed that control cell lines consistently exhibited similar levels of *panNRXN1* and its main isoforms, despite genetic background variability. Interestingly, the intron 5 deletion line 110_NXF show significant downregulation at day 0 (iPSC) (Two-way ANOVA: *padj = 0.0291, 95% CI) compared to controls but significantly increases at time points day 2 (Two-way ANOVA: *padj = 0.0375, 95% CI) and day 6 (Two-way ANOVA: ****padj = <0.0001, 95% CI). Conversely, cell line 109_NXM (intron 5 deletion) showed significantly reduced *panNRXN1* expression compared with controls at days 6 (Two-way ANOVA: ***padj = 0.0003, 95% CI) and 9 (Two-way ANOVA: **padj = 0.0040, 95% CI). Similarly, the cell line 092_NXF is significantly downregulated relative to controls at time points day 6 (Two-way ANOVA: **padj = 0.0012, 95% CI) and day 9 (Two-way ANOVA: *padj = 0.0109, 95% CI). The cell line 110_NXF demonstrated a significant increase in expression compared to 109_NXM on days 2 (Two-way ANOVA: ***padj = 0.0003, 95% CI), day 6 (Two-way ANOVA: ****padj < 0.0001, 95% CI), and day 9 (Two-way ANOVA: *padj = 0.0321, 95% CI). Additionally, compared to cell line 092_NXF, 110_NXF showed significantly higher expression levels on day 2 (Two-way ANOVA: **padj = 0.0027, 95% CI) and day 6 (Two-way ANOVA: ****padj < 0.0001, 95% CI) (**Figure 2C**).

**Figure 2:**
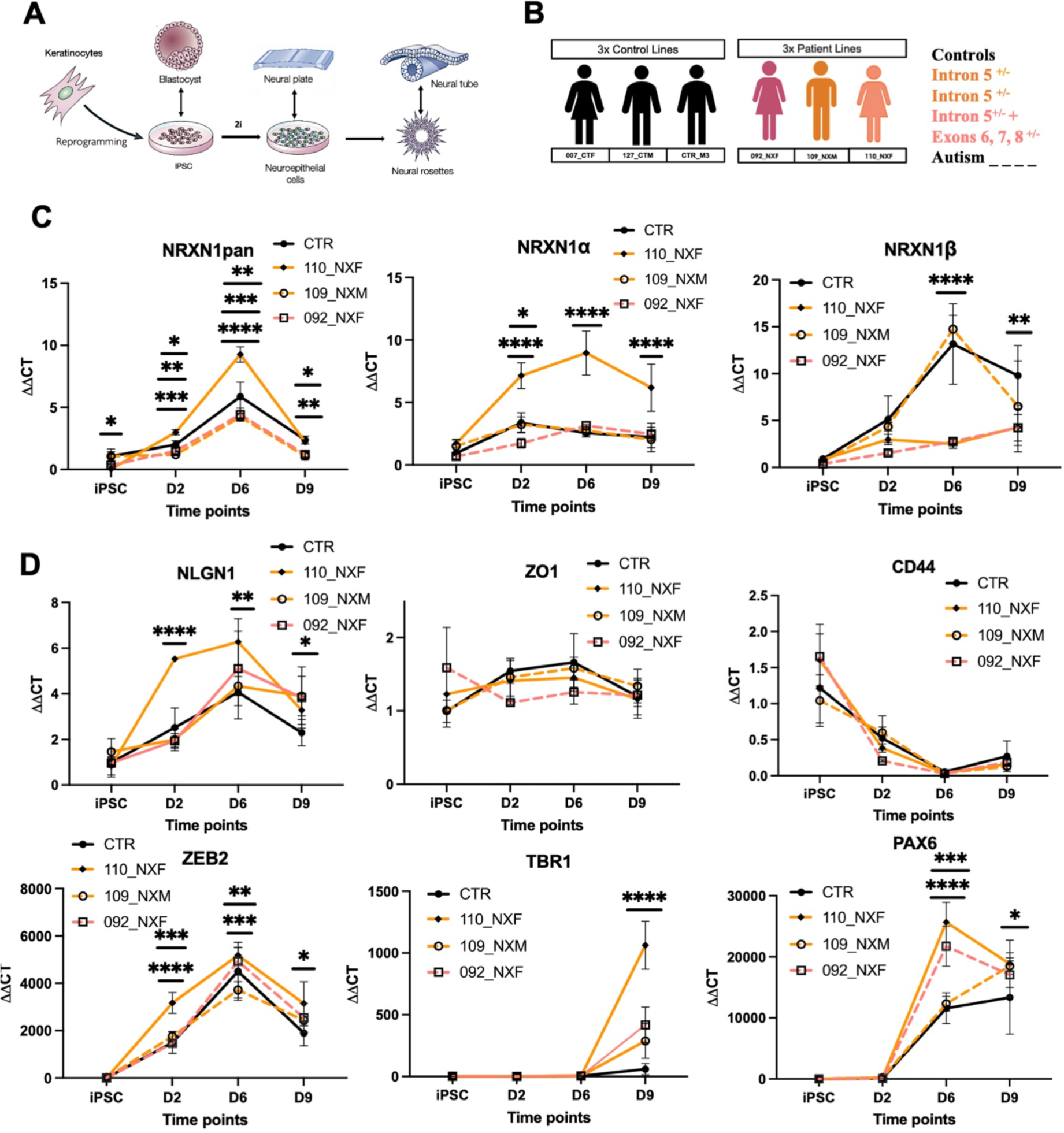
Gene expression patterns of *panNRXN1*, *NRXN1*, *NRXN1β*, *NLGN1*, ZO1*1*, *CD44*, *ZEB2*, *TBR1* and *PAX6* during early neural induction. **(A)** Schematic representation of the differentiation stages from human induced pluripotent stem cells (hiPSCs) to day 9 neural rosettes, outlining key developmental transitions. **(B)** Depiction of the genetic backgrounds in the control (black) and patient-derived cell lines, including the mother and son lines 110_NXF and 109_NXM with an identical deletion in *NRXN1* intron 5 (orange), and the unrelated line 092_NXF, which carries a deletion extending from intron 5 through exons 6, 7, and 8 (pink). **(C) I**llustrates the expression dynamics of *panNRXN1*, *NRXN1α*, and *NRXN1β* across early neuronal development, with peak expressions noted at day 6. *PanNRXN1* shows a significant downregulation in the 110_CTF cell line at the iPSC stage compared to controls, followed by significant upregulation on days 2 and 6. 109_NXM exhibits substantial downregulation on days 6 and 9 compared to controls. 092_NXF is similarly downregulated at days 6 and 9 relative to controls. 110_NXF shows pronounced upregulation compared to 109_NXM (days 2, 6, and 9) and 092_NXF (days 2 and 6). *NRXN1α* is markedly upregulated in 110_NXF across days 2, 6, and 9 compared to both controls and other deletion lines, while its expression is significantly reduced in 092_NXF compared with controls at day 2. *NRXN1β* expression is significantly reduced in the female deletion lines 110_NXF and 092_NXF at day 6 compared to controls and at day 9 relative to 110_NXF and 092_NXF. Whereas 109 NXM expression is upregulated at day 6 compared to 110_NXF and 092_NXF. **(D)** Expression Profiles of Other Relevant Neurodevelopmental Markers: *NLGN1* exhibits a significant increase in 110_NXF (days 2 and 6) compared to controls, and relative to 109_NXM and 092_NXF (day 2).109_NXM shows a significant increase in expression on day 9 compared to controls. *ZO1* levels increase on day 6, while *CD44* is consistently downregulated; however, these changes are not significant between cell lines. *PAX6* is significantly upregulated in 110_CTF (day 6 and 9) and 092_NXF (day 6) compared to controls. 109_NXM displays significant downregulation on day 6 compared to female lines 110_NXF and 092_NXF. *ZEB2* is upregulated in 110_NXF (days 2 and 9) relative to controls and compared to 109_NXM (day 2) and 092_NXF (day 6). 109_NXM shows upregulation compared to controls and downregulation compared to 092_NXF on day 6. TBR1 expression increases significantly on day 9 in all three NRXN1 deletion lines compared to controls, with 110_NXF also showing significant upregulation compared to 109_CTM and 092_NXF.

*NRXN1α* expression was significantly upregulated in the *NRXN1* deletion line 110_NXF at days 2 (Two-way ANOVA: ****padj < 0.0001, 95% CI), day 6 (Two-way ANOVA: **padj = 0.0012, 95% CI) and day 9 (Two-way ANOVA: **padj = 0.0012, 95% CI) following neuronal induction. This upregulation is also significant when compared with cell lines 109_NXM and 092_NXF at days 2, 6, and 9 (Two-way ANOVA: ****padj < 0.0001). Whereas the expression in cell line 092_NXF is significantly downregulated compared to controls on day 2 (Two-way ANOVA: *padj = 0.0468, 95% CI) (**Figure 2C**). Analysis of the *NRXN1β* isoform reveals significant downregulation in the female *NRXN1* deletion cell lines 110_NXF and 092_NXF compared to controls, with statistically significant reductions observed at day 6 (Two-way ANOVA: ****padj < 0.0001, 95% CI for both lines). Further declines in expression continue at day 9, with 110_NXF and 092_NXF showing continued significant downregulation relative to controls (Two-way ANOVA: **padj = 0.0097 and **padj < 0.0087, 95% CI, respectively). Additionally, the cell line 109 NXM was significantly upregulated compared with the cell line 110_CTF (Two-way ANOVA: ****padj = <0.0001, 95% CI) and 092_NXF (Two-way ANOVA: ****padj = <0.0001, 95% CI) at day 6. (**Figure 2C**). These data suggest that deletion of intron 5 of *NRXN1* results in altered isoform expression during early corticogenesis.

Previous studies have demonstrated that bi-allelic exonic deletions in are associated with altered expression of NRXN1 signalling molecules as well as cell fate commitment (Lam et al., 2019; Sebastian et al., 2023). Therefore, to understand whether similar alterations could be observed in our *NRXN1* deletion lines that are characterised by intronic deletions, we assessed the expression of key genes associated with NRXN1 signalling and cell fate markers. *NRXN1* expression was also associated with altering the expression of its binding partner *NLGN1*, key neurodevelopmental markers (*PAX6*, *ZEB2*, *TBR1*), and the tight junction marker *ZO1*, which is highly expressed at the inner lumen of neural rosettes (**Figure 2.D**). Our findings indicate that expression of *NLGN1* was high during early development, particularly at days 2 and 6 following 2i neural induction. While expression patterns remained consistent across different cell lines, *NLGN1* transcript levels were significantly higher in the *NRXN1* deletion line 110_NXF on day 2 (Two-way ANOVA: ****padj = <0.0001, 95% CI) and day 6 (Two-way ANOVA: **padj = 0.0029, 95% CI) of differentiation compared with controls. Additionally, on day 2, the increase in *NLGN1* is significantly more pronounced in 110_NXF when compared with cell lines 109_NXM (Two-way ANOVA: ****padj < 0.0001, 95% CI) and 092_NXF (Two-way ANOVA: ****padj = 0.0001, 95% CI). The cell line 109_NXM shows significant increase of expression at day 9 (Two-way ANOVA: *padj = 0.0438) compared with controls (**Figure 2D**).

*ZO1* transcript levels increased on day 6, remaining consistent and not differing across all hiPSC lines. (**Figure 2D**). As previous studies observed increased expression of astroglia cells in hiPSCs with bi-allelic exonic *NRXN1* deletions, we assessed the expression of the glial fate marker *CD44*. This gene showed reduced expression from day 2 in all differentiating cell lines (**Figure 2D**), suggesting that neither control nor *NRXN1* deletion lines were adopting a glial fate. Interestingly, the neurodevelopmental marker *PAX6* showed significantly elevated expression in the female *NRXN1* deletions lines 110_NXF (Two-way ANOVA: ****padj = <0.0001, 95% CI) and 092_NXF (Two-way ANOVA: *****padj = <0.0001, 95% CI) compared to controls at day 6 and at day 9 only for the cell line 110_NXF (Two-way ANOVA: *padj = <0.0114, 95% CI). While the cell line 109_NXM is significantly downregulated compared to the female lines 110_NXF (Two-way ANOVA: ****padj = <0.0001, 95% CI) and 092_NXF (Two-way ANOVA: ***padj = 0.0007, 95% CI) at day 6. *ZEB2*, a marker of neuroepithelial differentiation, showed elevated expression in the 110_NXF (Two-way ANOVA: ****padj = <0.0001, 95% CI) line at day 2 and day 9 (Two-way ANOVA: *padj = <0.0114, 95% CI) compared to control lines. Similarly, the deletion line 109_NXF at day 6 (Two-way ANOVA: ***padj = 0.0001, 95% CI) shows significantly elevated expression of *ZEB2* transcripts compared to controls. While the cell line 110_NXF is significantly upregulated at day 2 compared with 109_NXM (Two-way ANOVA: ***padj = 0.0005, 95% CI) and 092_NXF (Two-way ANOVA: ****padj = <0.0001, 95% CI) and at day 6 only compared to the cell line 109_NXM (Two-way ANOVA: ***padj = 0.0003, 95% CI). The line 109_NXM is significantly downregulated compared with 092_NXF at day 6 (Two-way ANOVA: **padj = 0.0030, 95% CI). The forebrain marker *TBR1* showed a significant increase in all *NRXN1* deletion lines relative to controls (Two-way ANOVA: ****padj = <0.0001, 95% CI). *TBR1* transcripts were also elevated in the cell line 110_NXF compared to the cell lines 109_CTM (Two-way ANOVA: ****padj = 0.0001, 95% CI) and 092_NXF (Two-way ANOVA: ****padj = <0.0001, 95% CI) at day 9, highlighting a potentially altered developmental trajectory influenced by altered *NRXN1* isoform expression levels. Collectively, these findings reveal that deletions in intron 5 are associated with altered expression of *NRXN1* main isoforms and the genes *NLGN1, ZEB2, TBR1* and *PAX6* across cortical differentiation, but that variations across NRXN1 deletions lines were evident suggesting an influence of genetic background or biological sex.

### Intronic *NRXN1* deletion lines exhibit altered isoform expression in immature neurons

After 30 days of neural induction, we observed the formation of immature neurons across all cell lines. At this mature stage, we systematically assessed the expression of *NRXN1* isoforms and several key developmental markers within three control and three *NRXN1* deletion lines (**Figure 3A-B**). Notably, while the expression of *panNRXN1* and its main isoforms, *NRXN1α* and *NRXN1β*, increased from earlier developmental stages, *panNRXN1* expression was significantly reduced in all *NRXN1* deletion lines: 110_CFT (One-way ANOVA: ***padj = 0.0004, 95% CI), 109_NCM (One-way ANOVA: ****padj = <0.0001, 95% CI) and 092_NXF (One-way ANOVA: **padj = 0.0059, 95% CI) compared to controls. Additionally, the deletion line 109_NXM show significant reduced expression of *panNRXN1* compared with 092_NXF (One-way ANOVA: *padj = 0.0118, 95% CI) (**Figure 3C**). In contrast, *NRXN1α* maintained a consistent expression profile across all cell lines. However, *NRNX1β* expression was reduced in lines with intron 5 deletions and was significantly downregulated in the *NRXN1* deletion line 109_NXM line (One-way ANOVA: **padj = 0.0074, 95% CI) relative to controls (**Figure 3C**). By day 30, the levels of *NLGN1*, ZO*1*, and the neuronal markers *PAX6* and *ZEB2* were similarly expressed across all immature neurons, indicating a uniform developmental progression (**Figure 3D**). The expression of the forebrain marker *TBR1* did not differ between control and *NRXN1* deletion lines. Notably, *CD44* transcript levels were significantly upregulated only in the female *NRXN1* deletion line 110_NXF (One-way ANOVA: *padj = 0.0150, 95% CI) at day 30 compared with controls. Additionally, significant increase of expression was observed in the cell line 110_CTF compared to the lines 109_NXM (One-way ANOVA: *padj = 0.0120, 95% CI) and 092_NXF (One-way ANOVA: *padj = 0.0153, 95% CI) at this developmental stage (**Figure 3.D**). Collectively, these findings reveal that cell lines exhibited similar expression patterns for fate markers. However, *NRXN1* isoform expression was altered in *NRXN1* deletion lines compared to control lines, with observed variability likely influenced by genetic background differences or biological sex between deletion lines.

**Figure 3:**
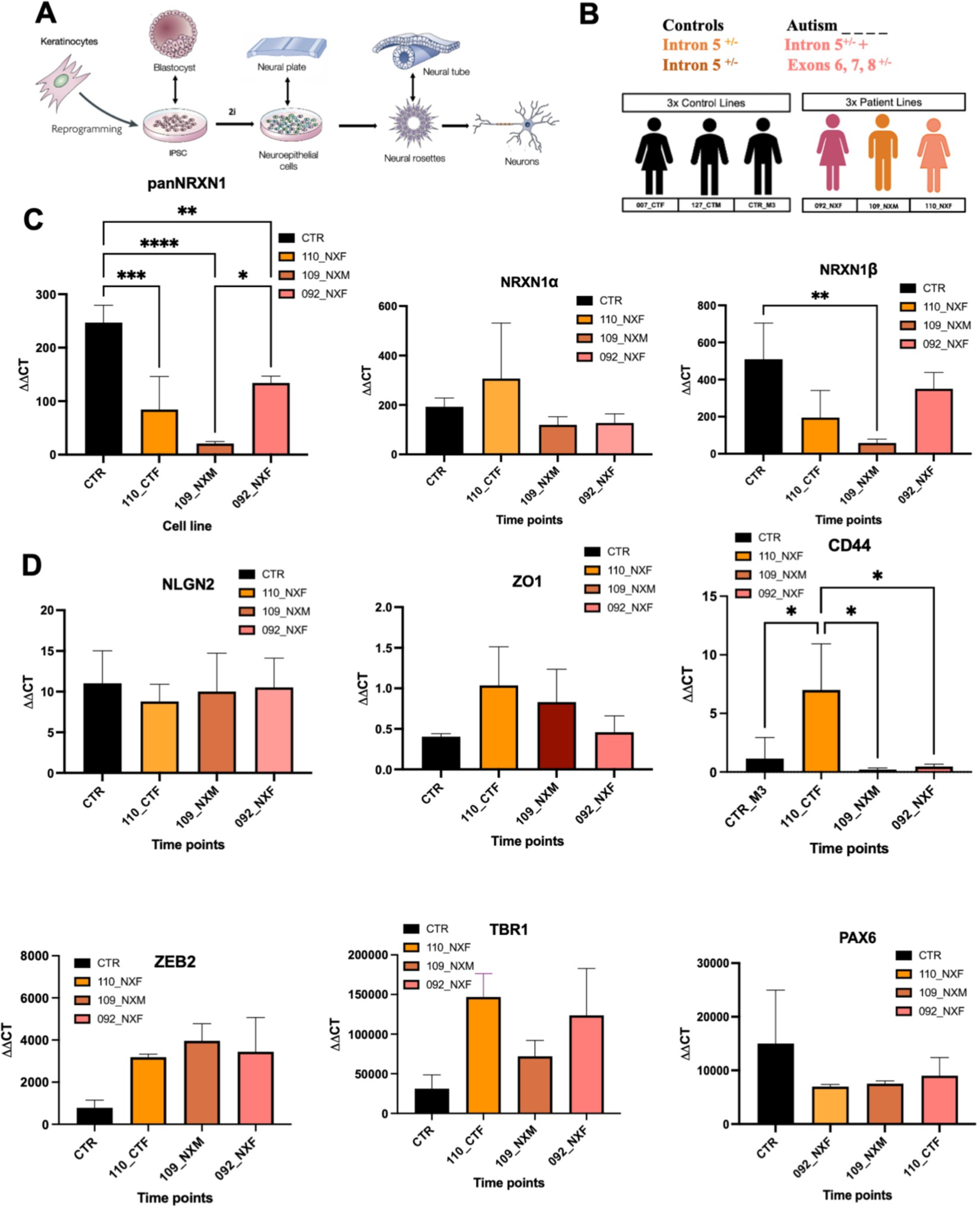
Gene expression patterns of *panNRXN1*, *NRXN1α*, *NRXN1β*, *NLGN1*, ZO1*1*, *CD44*, *ZEB2*, *TBR1* and *PAX6* in immature neurons. **(A)** Schematic illustration of the differentiation stages from human induced pluripotent stem cells (hiPSCs) to day 30 immature neurons, highlighting key developmental transitions. **(B)** Overview of genetic backgrounds in control deletion cell lines. **(C)** Comparative expression analysis at day 30 of neuronal differentiation: Immature neurons from NRXN1 intron 5 deletion lines 110_CFT, 109_NCM and 092_NXF exhibit a significant reduction in pan NRXN1 expression compared to controls. Additionally, differential expression was detected between the cell lines 109_NXM and 092_NXF. NRXN1α maintains consistent expression levels across all cell lines, whereas NRXN1β is significantly downregulated in the 109_NXM relative to controls. **(D)** Levels of *NLGN2*, ZO*1*, *ZEB2*, *TBR1*, and *PAX6* remain consistent between control and *NRXN1* deletion lines, indicating no major differences in the expression of these developmental markers. The expression of the glial marker *CD44* is significantly elevated in the 110_NXF cell line at day 30 compared with controls and the other two deletion lines the deletion lines 109_NXM and 092_NXF.

### Molecular characterization and enrichment of rare variants in *NRXN1* deletion cell lines compared to controls

To gain an insight into the molecular differences between *NRXN1* deletion (092_NXF, 109_NXM, 110_NXF) and control (007_CTF, 127_CTM) lines, we performed a burden analysis comparing rare variant counts using whole genome sequencing (WGS) data generated from hiPSCs. The control group had 237 predicted deleterious variants in 233 genes across two samples. *NRXN1* deletion hiPSC lines showed 401 predicted deleterious variants in 395 genes across three samples. Welch two-sample t-tests revealed a significant increase in both gene count (p-value = 0.00039) and variant count (p-value = 0.00053) in the *NRXN1* deletion cohort compared to controls (**Figure 4A**), highlighting a substantial enrichment of rare variants in the *NRXN1* deletion lines.

**Figure 4:**
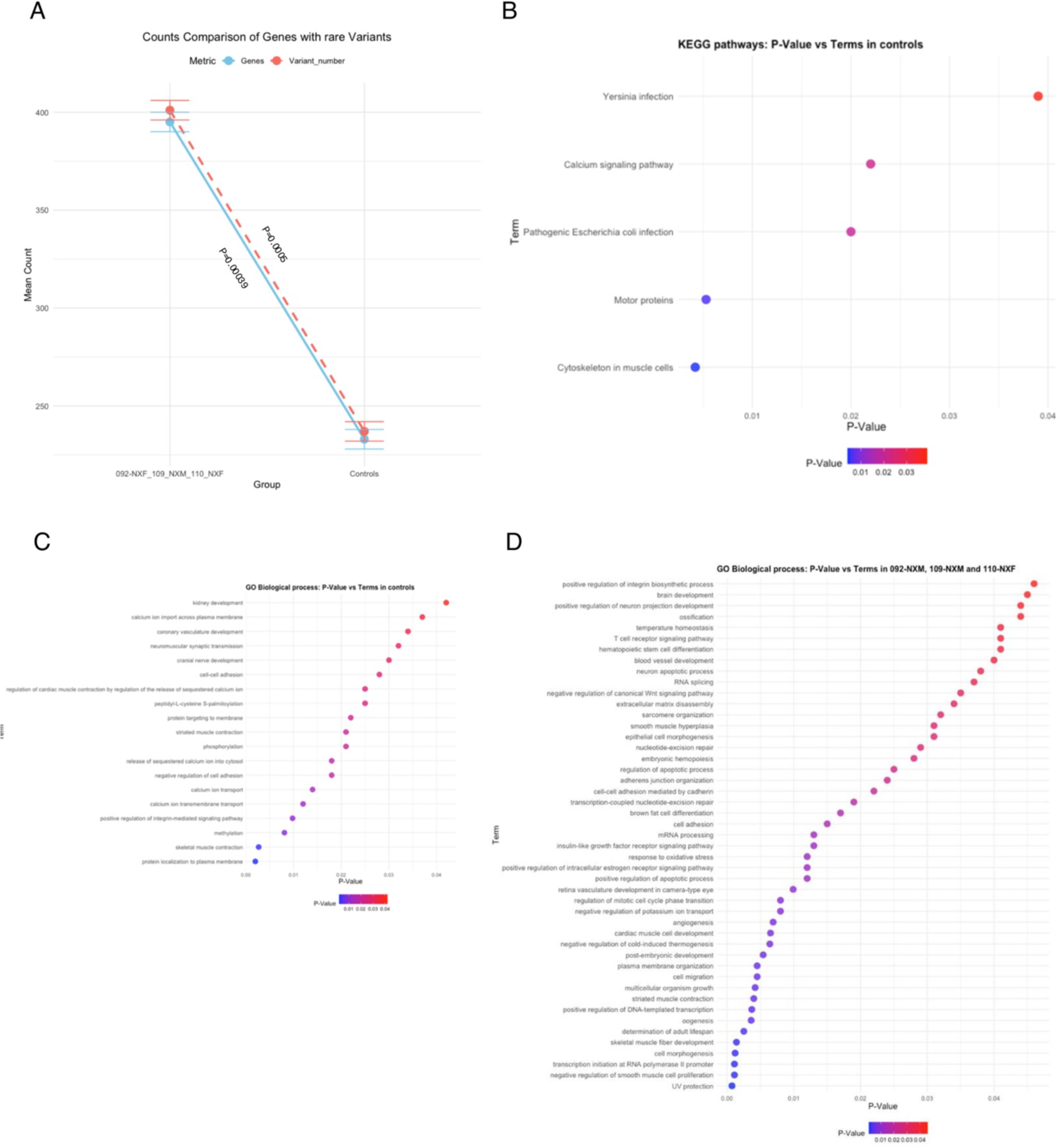
Analysis of predicted deleterious rare variants in *NRXN1* deletions and control lines using whole genome sequencing data. **A)** Statistical enrichment analysis comparing the number of genes and rare variant between *NRXN1* deletion cell lines and controls. **B)** Significant KEGG pathways enriched in control cell lines based on genes with rare variants. **C)** Enriched Gene Ontology (GO) biological processes in control cell lines, identified using genes with rare variants. **D)** Enriched GO biological processes in *NRXN1* deletion cell lines, identified using genes with rare variant.

Following burden assessment, we conducted a pathway enrichment analysis to explore the functional implications of the identified rare variants. In the control group, significant enrichment was observed in five KEGG pathways (p-value ≤ 0.05), including calcium signalling and motor proteins (**Figure 4B**). Gene Ontology Biological Processes (GO-BP) revealed enrichment in processes such as calcium transport, cell adhesion and methylation (**Figure 4C**). For cell lines with *NRXN1* deletions, we observed significant enrichment in processes related to brain development, neuron projection regulation, cell adhesion and RNA splicing (**Figure 4D**). No such enrichment was found in the control group.

We next conducted separate enrichment analyses for cell lines 109_NXM and 110_NXF to identify specific patterns within each line. Comparative enrichment analysis of genes with predicated deleterious variants from the 109_NXM and 110_NXF cell lines revealed enrichments in GO gene sets associated with post-embryonic development, cell morphogenesis, and cell adhesion in both hiPSC lines (**Figure 5A**). Examination of predicated deleterious variants in the 109_NXM cell line revealed significant enrichment of genes associated with neuron projection regulation, brain development and NMDA receptor clustering, along with gene expression regulation, which appeared specific to this hiPSC line (**Figure 5B**). Conversely, enrichment of genes associated with neuron development and cell morphogenesis appeared uniquely enriched in the 110_NXF line (**Figure 5C**). In summary, enrichment analysis reveals both common and unique findings for controls and *NRXN1* deletion cell lines. Both control and *NRXN1* deletion lines showed enrichment of genes associated with cell adhesion, but control lines had variants in genes related to calcium transport. *NRXN1* deletion cell lines, however, were uniquely enriched for processes such as brain development, neuron projection regulation, cell morphogenesis, and RNA processing. Moreover, *NRXN1* deletion lines showed unique enrichment for processes including those involved in regulation of gene expression and clustering of glutamate receptors.

**Figure 5:**
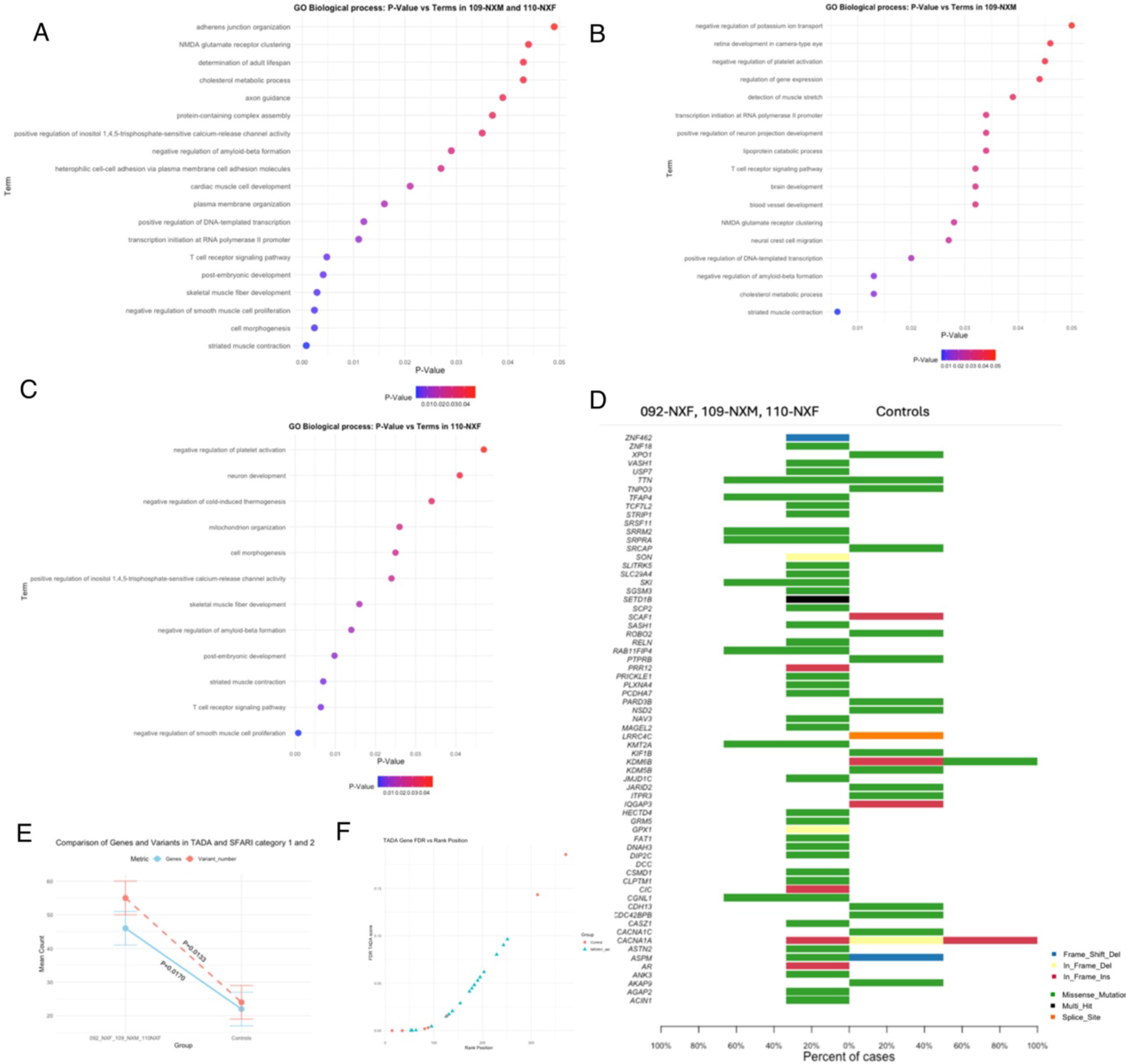
Analysis of autism associated rare and TADA genes in *NRXN1* deletions lines using whole genome sequencing. **A)** Gene Ontology (GO) biological pathways enriched using genes with rare variants from both the 109_NXM and 110_NXF cell lines. **B)** GO biological processes enriched using genes with rare variants exclusively from the 109_NXM cell line. **C)** GO biological processes enriched using genes with rare variants exclusively from the 110_NXF cell line. **D)** Bar plot illustrating the distribution of genes with rare variants from TADA and SFARI categories 1 and 2 across *NRXN1* deletion cell lines versus control cell lines. **E)** Statistical enrichment analysis comparing the number of TADA and SFARI categories 1 and 2 genes and variants in *NRXN1* deletion cell lines versus controls. **F)** relationship between the rank position and the False Discovery Rate (FDR) TADA score for the TADA genes found with moderate and rare variants. Each point on the scatter plot represents an individual gene, with its rank position on the x-axis and its TADA score on the y-axis.

To gain a deeper understanding of the genetic landscape and potential links with autism, we examined whether control or *NRXN1* deletions lines were enriched for rare variants in genes associated with autism categorized by the Transmission and De Novo Association (TADA) model (Sanders et al., 2015; Feliciano et al., 2019; Ruzzo et al., 2019; Satterstrom et al., 2020; Fu et al., 2022) and the SFARI database (categories 1 and 2) (Abrahams et al., 2013) In the control lines, we identified 24 predicted deleterious variants in 22 genes (**Figure 5D**). In contrast, the *NRXN1* deletion cell lines – 092_NXM, 109_NXM, and 110_NXF - exhibited a more complex genetic profile, with 55 variants in 46 genes (**Figure 5D**). Welch two-sample t-tests revealed a significant difference between the groups, with p-values of 0.017 for gene comparisons and 0.0133 for variant counts (**Figure 5E**), these results underscore an increased genetic burden of high-confidence autism genes in the *NRXN1* deletion cell lines.

### Intragenic *NRXN1* deletion hiPSC lines do not show altered cell fate or region identity

We next used RNA sequencing to explore the molecular mechanisms regulating neuronal formation and function in control cell lines and lines harbouring intronic deletions in *NRXN1*. We analysed neurons derived from hiPSCs at day 30, involving three control lines with one clone each and three *NRXN1* deletion lines with three clones each (**Figure 6A**). A sample distance plot and principal component analysis (PCA) revealed overall similarities among the samples, with no distinct clusters forming. Notably, clone 4 of cell line 109_NXM and clone 3 of 110_NXF exhibited slight segregation from the rest (**Figure 6B-C**). Furthermore, examination of the top 20 most differentially expressed genes across all samples indicated a conserved expression pattern between cell lines (**Figure 6D**).

**Figure 6:**
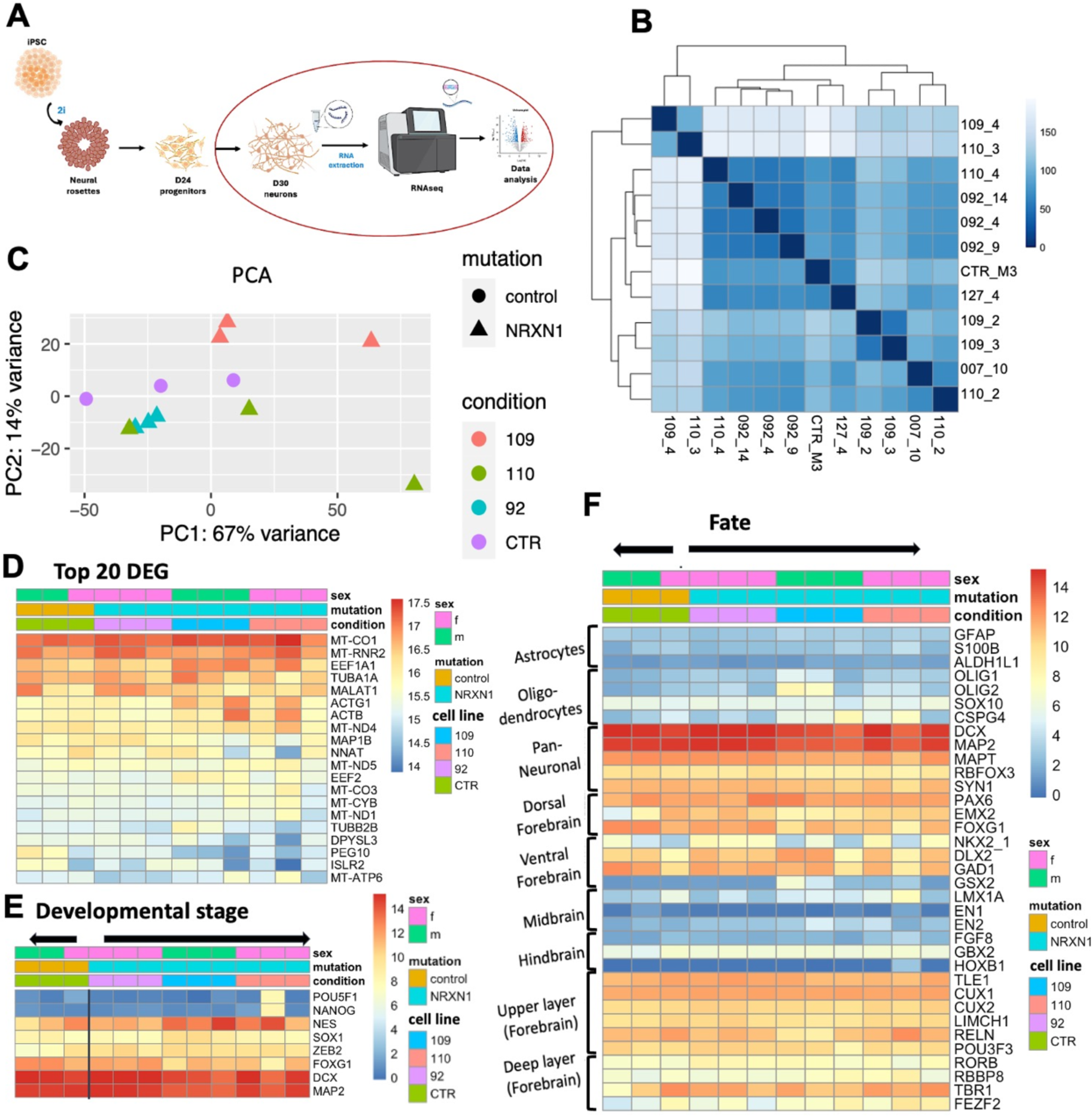
Gene expression patterns in immature neurons. **(A)** Schematic representation of the differentiation process from hiPSCs to day 30 neurons, including stages for RNA sequencing analysis. **(B-C)** Sample distance plot (B) and principal component analysis (PCA) (C) illustrate the overall similarity among samples, indicating no significant clustering by cell line type. **(D)** Visualization of the top 20 DEGs between control lines (green) and *NRXN1* deletion lines (light blue). The expression patterns from specific cell lines are shown: 109_NXM (pink), 110_NXF (red), and 092_NXF (blue). **(E)** Heatmap of gene expression across all cell lines, displaying low expression levels of pluripotency markers, moderate expression of neuroepithelial markers, and high expression of forebrain-specific markers. **(F)** Expression patterns of cell fate markers reveal low expression of astrocyte, oligodendrocyte, midbrain, and hindbrain markers, contrasting with high expression of markers associated with forebrain fate, including pan-neuronal, dorsal and ventral forebrain, and both upper and deep layer neurons.

To confirm that *NRXN1* deletions lines had a similar developmental trajectory and acquisition of cell fate compared to control lines following cortical differentiation, we assessed the expression of cell fate and developmental markers. As expected, day 30 hiPSC-derived neurons had negligible expression of pluripotency genes (*POU5F1*, *NANOG*), intermediate levels of neuroepithelial markers (*NES*, *SOX1*, *ZEB2*), and high expression of pan forebrain and neuronal genes (*FOXG1*, *DCX*, *MAP2*) in both control and *NRXN1* deletion lines, indicating similar developmental stages (**Figure 6E**). Moreover, we found low expression of genes associated with astrocytes, oligodendrocytes cell fates, or of non-forebrain regions such as the midbrain or hindbrain suggesting that the differentiation of all lines into cortical neuronal fates was similar across all cell lines (**Figure 6E**). Consistent with this, markers indicative of pan-neuronal, dorsal and ventral forebrain, and both upper and deep layer neurons were similarly expressed across all lines (**Figure 6F**). This demonstrates that both control and *NRXN1* deletion lines uniformly acquire a forebrain fate, underscoring a minimal impact of *NRXN1* intronic deletions on cell fate acquisition within the forebrain lineage (**Figure 6F**).

### Impact of *NRXN1* Intronic Deletions on Neuronal Gene Expression and Autism-Related Pathways

Previous research has demonstrated that exonic deletions in *NRXN1* can lead to significant alterations in molecular transcriptomic profiles (Flaherty et al., 2019; Lam et al., 2019; Sebastian et al., 2023). Building on this, we investigated whether intronic deletions in *NRXN1* are similarly associated with molecular changes and disruptions in regulatory pathways. We initially compared the gene expression profiles of control lines with all *NRXN1* deletion lines, identifying 54 differentially expressed genes (DEGs): 29 were significantly upregulated and 25 downregulated in the deletion lines compared to controls (**Figure 7A-B**) (**Supplementary table 1**). Given the strong association between molecular alterations in genes like *NRXN1* and autism (Onay et al., 2016; Ishizuka et al., 2020; Cooper et al., 2024) we sought to determine if there was an enrichment of DEGs between control and *NRXN1* deletion lines with the expression of autism-associated genes. We referenced the curated gene variants from the Simons Foundation Autism Research Initiative (SFARI) Gene database (Abrahams et al., 2013), specifically category 1 (high confidence genes) and category 2 (strong candidate genes). The analysis revealed no statistically significant overlap, with only the gene *SLC24A2* from SFARI category 2 showing significant reduction in the *NRXN1* deletion lines (**Figure 7C**) (**Supplementary table 2**).

**Figure 7:**
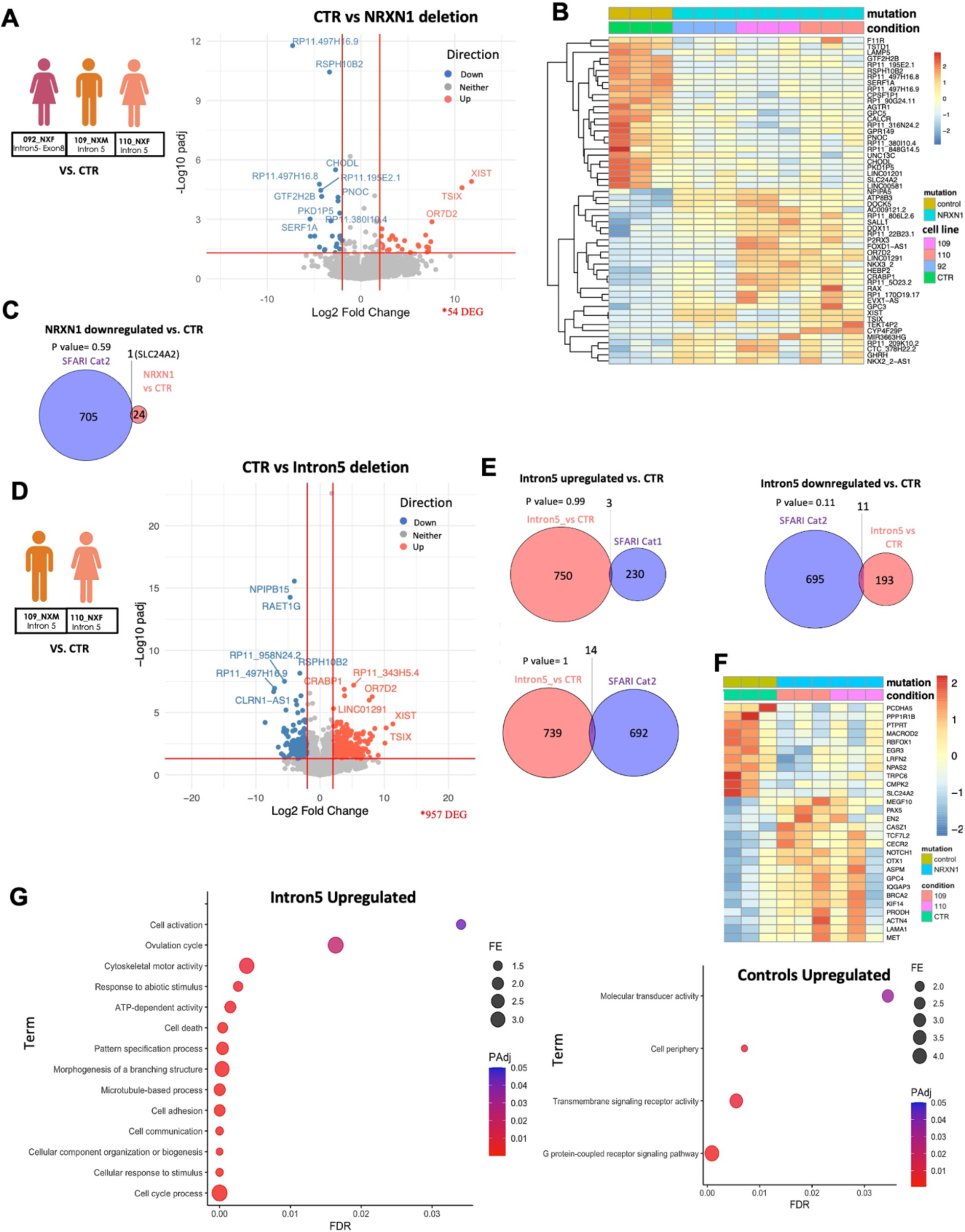
Differential Gene Expression and Pathway Analysis in *NRXN1* Deletion Lines. **(A)** DEG analysis comparing *NRXN1* deletion lines, including 092_NXF (pink), 109_NXM (orange), and 110_NXF (light orange), against control cell lines. A volcano plot illustrates 54 significant DEGs identified between these groups. **(B)** Heatmap displaying the upregulated and downregulated genes in the *NRXN1* deletion lines relative to control lines, highlighting the magnitude and direction of expression changes. **(C)** Visualization of the overlap between genes downregulated in the *NRXN1* deletion lines (pink) and the SFARI database category 2 list (lilac), identifying one overlapping gene. **(D)** Comparative DEG analysis between intron 5 deletion lines 109_NXM (orange) and 110_NXF (light orange), and control lines, revealing 957 DEGs. **(E)** Analysis of overlap between SFARI categories 1 and 2 with genes upregulated in intron 5 deletion lines, showing 3 and 14 genes respectively. Additionally, 11 genes downregulated in intron 5 deletion lines are also listed in SFARI category 2. **(F)** Heatmap illustrating the expression patterns of DEGs between intron 5 deletion lines and controls that are included in SFARI gene lists. **(G)** GO analysis plots demonstrating enriched biological pathways for genes upregulated and downregulated in intron 5 deletion lines compared to controls.

Further comparative analysis between the control lines and the cell lines sharing an identical deletion in intron 5 (110_NXF and 109_NXF), revealed 957 DEGs **(Supplementary table 1).** Among these, 753 genes were significantly upregulated and 204 downregulated. Of the upregulated genes, only 3 genes overlapped with SFARI category 1 and 14 in category 2 gene sets **(Supplementary table 2).** Additionally, 11 downregulated genes were also identified as overlapping with SFARI category 2 genes (**Figure 7D-F**). Gene Ontology (GO) analysis of the 753 upregulated genes highlighted enrichment in biological pathways related to cell adhesion, communication, and morphogenesis among others, suggesting these pathways might be affected in the *NRXN1* intron 5 deletion lines compared to controls. Conversely, pathways associated with molecular transducer and signalling receptor activity and others were enriched among the downregulated genes, indicating potential disruptions in these processes (**Figure 7G**) (**Supplementary table 3)**.

### Transcriptomic Differences and Synaptic Pathway Alterations in the *NRXN1* Deletion Line 109_NXM

We next conducted transcriptomic analysis comparing cell line 109_NXM, derived from an autistic male, with line 110_NXF, derived from his mother (no diagnosis): both cell lines carry identical deletions in intron 5 **(Supplementary table 1)**. This comparison revealed 271 differentially expressed genes (DEGs), with 58 genes significantly upregulated and 213 genes downregulated in the male line compared to the maternal line (**Figure 8A**). Among these, DEGs overlapping with the SFARI autism-related gene list include two from category 1 and three from category 2 in the upregulated set, while five from category 1 and eight from category 2 were found in the downregulated set (**Figures 8B-C**) (**Supplementary table 2)**. Significantly, genes downregulated in the male patient’s line were enriched for Gene Ontology (GO) terms related to synaptic function and development (**Figure 8D**) (**Supplementary table 3)**. Furthermore, analysis for overlap of DEGs with synaptic gene genes using SynGo (Koopmans et al., 2019), revealed significant enrichment for presynaptic, postsynaptic, and synaptic signalling pathways (**Figure 8E**).

**Figure 8:**
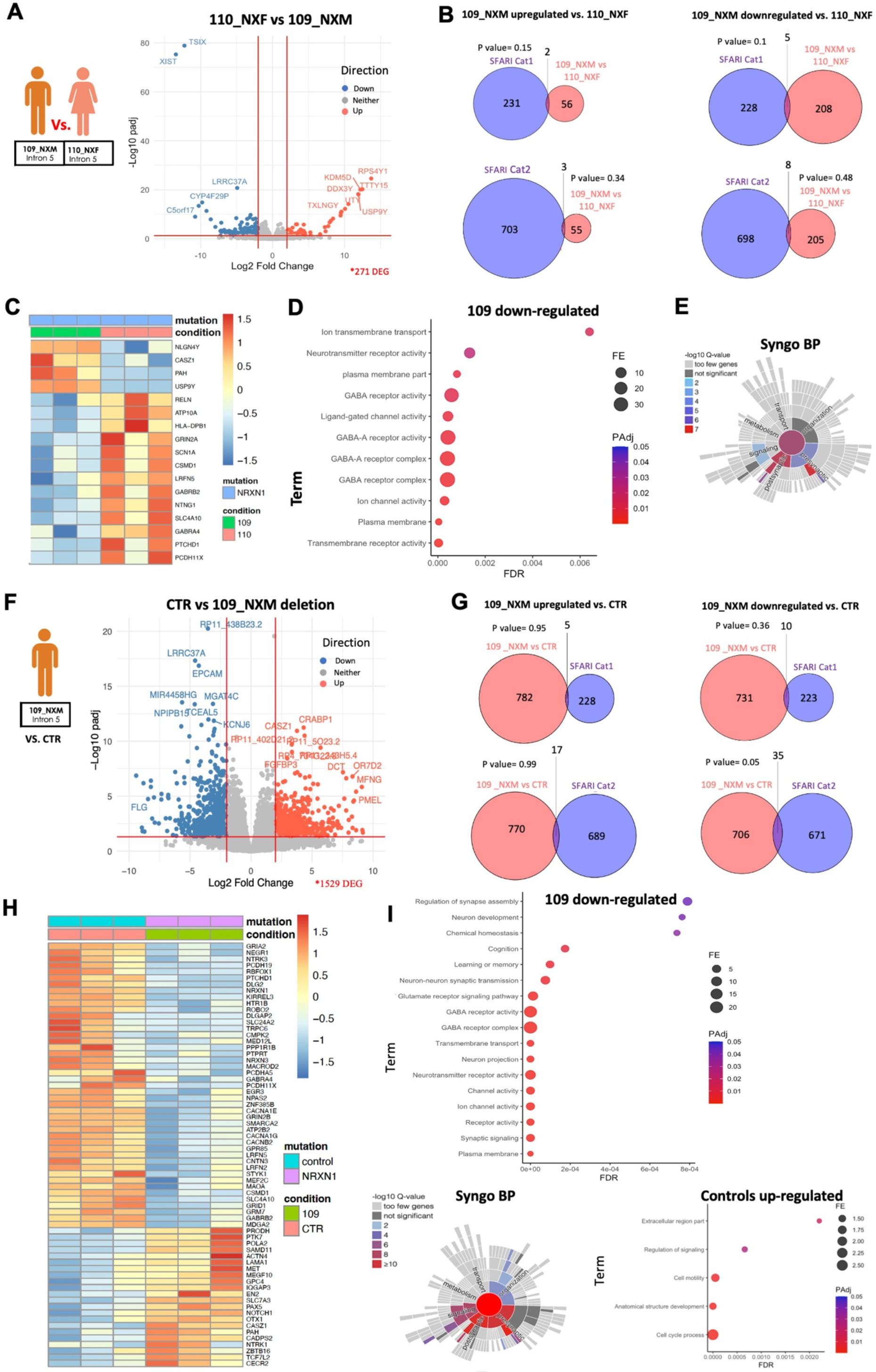
Differential Gene Expression and Pathway Analysis in *NRXN1* Deletion Line 109_NXM. **(A)** Differential expression analysis comparing *NRXN1* deletion line 109_NXM with the cell line 110_NXF derived from the participant’s mother. A total of 271 genes were identified as significantly differentially expressed between these lines. **(B)** Venn diagrams illustrating the overlap between genes upregulated and downregulated in cell line 109_NXM relative to line 110_NXF (pink), and their correspondence with genes listed in the SFARI database categories 1 and 2 (lilac). **(C)** Heatmap displaying the expression patterns of upregulated and downregulated genes in the *NRXN1* deletion line 109_NXM that correspond with SFARI gene lists. **(D)** Gene Ontology (GO) analysis plots showing enriched biological pathways for genes downregulated in *NRXN1* deletion line 109_NXM compared to line 110_NXF. **(E)** Schematic depiction of synaptic pathways significantly enriched among the downregulated genes in 109_NXM, based on SynGo ontology. **(F)** DEG analysis comparing cell line 109_NXM with control lines, identifying 1529 differentially expressed genes. **(G)** Diagrams illustrating the overlap between genes upregulated and downregulated in cell line 109_NXM compared to controls (pink) and their correspondence with the SFARI database categories 1 and 2 (lilac). **(H)** Heatmap displaying the expression patterns of upregulated and downregulated genes in the NRXN1 deletion line 109_NXM relative to controls that are included in the SFARI gene lists. **(I)** GO analysis plots revealing enriched biological pathways for genes both downregulated and upregulated in the NRXN1 deletion line 109_NXM compared to controls, highlighting the significant enrichment of synaptic pathways among the downregulated genes according to SynGo ontology.

When compared to control lines, the autistic male cell line 109_NXM showed 1529 DEGs (**Figure 8F**). These included 787 genes upregulated in the *NRXN1* deletion line, with overlaps of 5 genes with SFARI category 1 and 17 with category 2 **(Supplementary table 2)**. Furthermore, of then 741 DEGs showing reduced expression, 10 overlapped with SFARI category 1 and 35 with category 2 (**Figures 8G-H**). Notably, the 741 DEGs were enriched for biological functions regulating synapses and synaptic function, such as synaptic transmission, GABA receptor activity, channel activity, and synaptic signalling (**Figure 8I**) (**Supplementary table 3)**. These genes are significantly involved in synaptic organization including presynaptic and postsynaptic activities, as detailed in SynGo ontology (**Figure 8I**). Together, these findings indicate that alterations in synaptic formation and function, which have been previously linked to autism (Zoghbi et al., 2012; Kathuria et al., 2018; Adhya et al., 2021), are particularly present in the *NRXN1* deletion line derived from an autistic individual. Such alterations offer insights into the molecular mechanisms potentially underpinning the autism phenotype observed in the cell line.

### *NRXN1* deletion lines display altered dendrite outgrowth

Previous research has linked NRXNs to dendritogenesis in *Drosophila* and *Xenopus* neurodevelopment via NLGN1. NRXN1 binding partners NLGN3 and NLGN4X have also been linked to neuritogenesis in immature human neurons (Chen et al., 2010; Constance et al., 2018; Gatford et al., 2022). Given that *NRXN1* deletion lines showed altered *NRXN1* isoform expression, were enriched for rare variants associated with cell adhesion and neuronal projection regulation, and furthermore, immature neurons displayed altered expression of genes associated with cell adhesion, morphogenesis and synapse function, we hypothesized that patient lines may exhibit aberrant dendrite formation. HiPSCs from all lines were differentiated into cortical neurons until day 24, then fixed/stained for the dendrite marker MAP2 and DAPI, and were subjected to a high content screening analysis pipeline. This analysis pipeline was designed to automatically identify, trace, and quantify MAP2-positive ‘neurites’ in terms of ‘neurite number’, ‘neurite length’, and the number of branch points per neuron (Shum et al., 2020). Analysis of day 24 cortical neurons revealed a clear increased dendrite outgrowth phenotype in all *NRXN1* deletion lines compared to control lines (**Figure 9A**). Interestingly, differences in dendrite outgrowth could also be observed between *NRXN1* deletion lines. Averaging across all lines revealed that all patient lines exhibited significantly increased dendrite number (152.18%) compared to all control lines (‘neurite number’ per neuron CTR_M3, 1.58±0.12; 007_CTF, 0.95±0.02; 127_CTM, 0.91±0.15 – Averaged controls, 1.15; 092_NXF, 3.16±0.15; 109_NXM, 2.79±0.03; 110_NXF, 2.72±0.17 – Averaged patients, 2.89. One-way ANOVA: *F*(5,12)=70.57, *p*<0.0001, η^2^=0.97). Bonferroni post-hoc comparisons revealed the precise differences between individual control lines and individual patient lines, demonstrating that neurons generated from the 092_NXF line had the greatest number of dendrites per neuron (**Figure 9B**). *NRXN1* deletion lines also exhibited significantly increased dendrite length (156.52%) compared to all control lines (‘neurite length’ per neuron: CTR_M3, 75.68±7.98 µm^2^; 007_CTF, 54.93±1.98 µm^2^; 127_CTM, 35.27±1.52 µm^2^ – Averaged controls, 55.29 µm^2^; 092_NXF, 186.00±2.53 µm^2^; 109_NXM, 122.40±5.99 µm^2^; 110_NXF, 117.10±1.73 µm^2^ – Averaged patients, 141.83 µm^2^. One-way ANOVA: *F*(5,12)=156.30, *p*<0.0001, η^2^=0.99). Once more, Bonferroni post-hoc comparisons revealed the precise differences between individual control lines and individual patient lines, with 092_NXF *NRXN1* deletion line exhibiting significantly increased dendrite length compared to all other lines (**Figure 9C**). Similarly, *NRXN1* deletion lines also exhibited significantly increased branching (188.37%) compared to all control lines (branch point number per neuron: CTR_M3, 0.76±0.07; 007_CTF, 0.35±0.03; 127_CTM, 0.19±0.01 – Averaged controls, 0.43; 092_NXF, 1.78±0.11; 109_NXM, 1.01±0.08; 110_NXF, 0.94±0.03 – Averaged patients, 1.24. One-way ANOVA: *F*(5,12)=79.02, *p*<0.0001, η^2^=0.97). *NRXN1* deletion line 092_NXF had increased branch number compared to all other lines as determined by Bonferroni post-hoc comparisons (**Figure 9D**). Collectively, these data indicate that immature neurons differentiated from hiPSCs lines carrying intronic *NRXN1* deletions generate neurons with larger dendritic morphologies.

**Figure 9:**
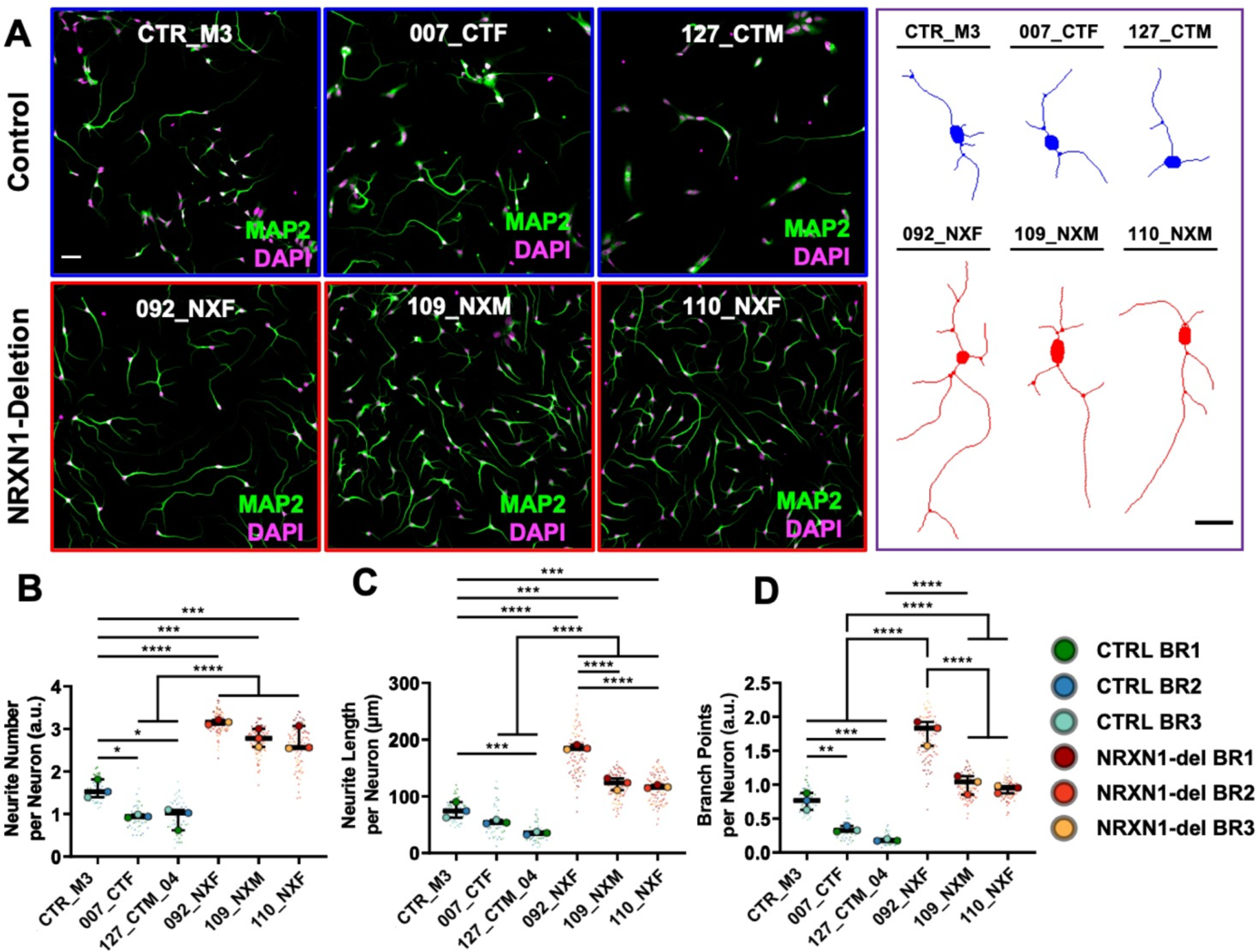
Intragenic *NRXN1* deletions are associated with increased dendrite outgrowth. (A) Representative images showing day 24 human induced pluripotent stem cell (hiPSC)-derived cortical neurons from control and *NRXN1* deletion lines and representative individual neurons extracted from the high-content screening dataset illustrating increases in MAP2-positive dendrite number, length, and branch point number in all control and *NRXN1* deletion lines. Scale bar = 100 µm. (B) Combined box plot and violin plot that showing *NRXN1* deletion lines have significantly increases dendrite number in hiPSC-derived neurons compared to hiPSC-derived neurons. (C) Plot data showing intronic *NRXN1* deletions associate with increased dendrite length in hiPSC-derived neurons. (D) Plot data showing intronic *NRXN1* deletions significantly increases branch point number in hiPSC-derived neurons generated from all *NRXN1* deletion lines compared to hiPSC-derived neurons generated from control lines.

## Discussion

Although *NRXN1* exonic deletions have been well established to induce alterations in the development, gene expression profile, morphology and function of immature neurons (Pak et al., 2015; Flaherty et al., 2019; Lam et al., 2019; Sebastian et al., 2023) the impact of intronic deletions on molecular and cellular phenotypes remains less clear. Existing research, such as that by Lowther et al. (2017), indicates that not all intronic *NRXN1* deletions are benign. In this study, we have explored the molecular and cellular impacts of intron 5 deletions, focusing on the expression of major *NRXN1* isoforms and the molecular pathways underlying neuronal development. Using three control cell lines and three cell lines with overlapping *NRXN1* intron 5 deletions, we find that *NRXN1* deletion lines exhibit altered *NRXN1* isoform expression, suggesting that loss of intron 5 results in the dysregulation of isoform expression. WGS analysis of *NRXN1* deletion lines revealed significant enrichment for rare variants enriched for processes such as brain development, neuron projection regulation, as well as cell adhesion. These variants were also strongly associated with high-confidence autism genes, supported by robust statistical analysis. RNA Sequencing of immature neurons further revealed significant changes in genes encoding cell adhesion molecules as well as those involved in the development of morphology and synaptic function. Consistent with our molecular findings, neurons generated from *NRXN1* deletion lines exhibited larger dendritic morphologies, potentially indicating that *NRXN1* isoforms may be involved in regulating dendritogenesis. Taken together, these results demonstrate that intronic deletions in *NRXN1* influence the molecular and cellular landscape during neurodevelopment, highlighting the need to further study and consider the potential association of intronic deletions with neurodevelopmental and psychiatric conditions.

Based on previous studies demonstrating that exonic deletions in *NRXN1* result in altered isoform expression (Flaherty et al., 2019; Sebastian et al., 2023), we assessed the impact of intronic deletions on the expression of major *NRXN1* isoforms across different neurodevelopmental stages. Consistent with post-mortem studies (Jenkins et al., 2014) as well as hiPSC studies (Flaherty et al., 2019; Lam et al., 2019; Sebastian et al., 2023), we found that *NRXN1α* and *NRXN1β* increased in expression during the first 30 days of cortical differentiation, with a peak expression at day 6 and a larger peak at day 30. This observation reinforces the evidence that *NRXN1* is crucial during early neuronal development (Lam et al., 2019; Adhya et al., 2021; Sebastian et al., 2023). Of note, the expression of *NRXN1β* was consistently higher across most time points compared to *NRXN1α*, aligning with previous observations from human embryonic post-mortem tissue (Jenkins et al., 2014; Harkin et al., 2017). Deletions within intron 5 of *NRXN1* resulted in distinct expression profiles of *NRXN1α* and *NRXN1β* isoforms across cortical differentiation. While the *NRXN1* deletion in intron 5 does not disrupt the putative promoters or canonical splicing sites of *NRXN1α* and *NRXN1β*, the deletion may affect the sequences of alternative promoters, enhancers, or splicing regulators (Currant et al., 2013; Lowther et al., 2017). Such disruptions could be linked to the significant alterations observed in the expression patterns of *NRXN1* isoforms within *NRXN1* deletion lines. Interestingly, we observed significant differences in isoform expression between *NRXN1* deletion lines, even between 2 cell lines carrying identical intronic deletions derived from a familial paring (mother/son). It is likely that these differences are reflective of the diversity in genetic background between individuals. Indeed, while our analysis of WGS for predicted deleterious rare gene variants in genes associated with RNA splicing and other processes in all three patient lines, significant differences in variants were also observed between *NRXN1* deletion lines associated in processes such as gene regulation and NMDA receptor clustering, highlighting the inter-individual genetic differences. Many hiPSC studies attempt to account for such differences by increasing the number of donor lines used to provide an accurate understanding of common biological differences between case and control lines (Anderson et al., 2021; Dutan Polit et al., 2023). The findings in the current study demonstrate how taking into account inter-individual genetic variations may offer insight into differences in molecular and cellular phenotypes within specific sample groups.

In immature neurons, *NRXN1* deletion lines exhibited modest differences in gene expression between control and control and *NRXN1* deletion lines. Focused comparisons using only *NRXN1* intron 5 deletion lines revealed changes in the expression of genes critical to synaptic mechanisms, including cell adhesion, microtubule processes, and cytoskeletal activity, some of which are highly associated with autism (Parato & Bartolini, 2021; Jiang et al., 2022). Alterations in these pathways could disrupt cytoskeletal organization and dendrite elongation and branching, as observed in the *NRXN1* deletion lines (Miller & Suter, 2018). Comparison between lines harbouring the same intron 5 deletion revealed that the line 109_NXM exhibited alterations in crucial synaptic pathways, including neurotransmitter receptor activity, ion channel activity, and GABA receptor activity. These findings are in line with previous research highlighting synaptic impairments associated with *NRXN1* deletions (Pak et al., 2015; Flaherty et al., 2019; Lam et al., 2019; Sebastian et al., 2023). The interaction of the *NRXN1* intron 5 deletion with other genetic variants or sex-linked factors in the 110_NXM line may cumulatively alter its developmental trajectory, an effect not observed in the maternal line. Of note, the 109_NXM line was uniquely enriched with variants in genes associated with the regulation of NMDAR receptor clustering, which may contribute to some of the observed differences in alteration in synaptic pathways. These findings further underscore the potential influence of unique genetic architectures on the molecular landscape of *NRXN1* intronic deletion lines.

Consistent with predictions made from RNA sequencing data, neurons generated from *NRXN1* deletion lines displayed increased dendritic arborization compared to control neurons. While this enhanced dendritic outgrowth may increase dendritic field coverage, it could also reflect alterations in synaptic connectivity and neural network integration (Srivastava et al., 2012; Dong et al., 2015; Martínez-Cerdeño, 2017; Lanoue & Cooper, 2019; Palavalli et al., 2021). Previous work has demonstrated that interactions between *NLGN1* and *NRXN* are critical for filopodia stabilization and dendritogenesis in Xenopus and Drosophila (Chen et al., 2010; Constance et al., 2018). However, the role of *NRXN1* in neuritogenesis or dendritogenesis remains less clear. Contrasting studies show no effect of *NRXN1* deletions on dendritic arborization in ESC-derived neurons (Pak et al., 2018), while iPSC-derived neuron studies report decreased neurite branching from *NRXN1* exonic deletions (Flaherty et al., 2019). Such differences could be due to the nature and location of *NRXN1* deletion, or morphological timepoint investigated (i.e. neurite vs. dendrite). Molecular and cellular data generated in this study indicate that *NRXN1* intronic deletions disrupt key molecular players, such as adhesion molecules, which are key regulators of dendrite structure and formation (Srivastava et al., 2012; Gatford et al., 2022). In line with our molecular studies, we observed distinct morphological differences between *NRXN1* deletion lines. Interestingly, *NRXN1* deletion lines with identical deletions within intron 5 exhibited similar morphological profiles. In contrast, the deletion line derived from an unrelated individual displayed significantly larger dendritic arborisation. These differences may be due to the fact that this hiPSC line harbours a more extensive deletion, spanning from intron 5 to exon 8, in *NRXN1* or due to other variations in the genetic background of the line. Nevertheless, the robust alterations in dendritic morphology observed in both intron 5 specific deletion lines suggest that the molecular changes induced by this intragenic deletion may lead to divergent development of dendritic field coverage, which could influence synapse formation in developing neurons.

This study presents several limitations that should be considered when interpreting the results. Firstly, the small sample size, limited to only three controls and three *NRXN1* deletion lines, may reduce the statistical power and generalizability of the findings (Dutan Polit et al., 2023). Additionally, the analysis focused solely on major isoforms of *NRXN1*, overlooking the roles of multiple other isoforms – further studies using long-read technologies will be needed to overcome this limitation. Dendrite outgrowth was examined at a single time point, which may not capture the dynamic changes over time. The inherent variability in the genetic background of the cells may mask the effects of the *NRXN1* deletion mutation. Furthermore, transcriptomic analyses were not stratified by sex, which could obscure sex-specific differences in gene expression and the impact of the intronic deletion.

*NRXN1* deletions exhibit incomplete penetrance for clinical phenotypes. It has been suggested that the presence of co-occurring genetic factors may play a role in modulating this penetrance (Woodbury-Smith et al., 2017; Kasem et al., 2018; Cameli et al., 2021). Our research builds on this foundation by highlighting the impact of the different genetic landscapes within iPSC-derived neurons on the consequences of *NRXN1* intronic deletions. We hypothesize that variations in gene expression patterns and cellular phenotypes among cell lines with *NRXN1* deletions are directly influenced by the unique genetic makeup or sex-chromosome effects of each cell line. For instance, cell lines 110_NXF and 109_NXM, carrying the same *NRXN1* deletions in intron 5 exhibit markedly different expression patterns of *NRXN1* main isoforms, burden of predicted deleterious variants in high confidence autism genes, and molecular pathways associated with synaptic regulation and function. This suggests a complex aetiology that extends beyond a single genetic variant like *NRXN1*. Our findings align with research indicating a heightened male susceptibility to autism, potentially linked to male-biased genes regulating chromatin and immune functions, both recognized contributors to autism (Werling et al., 2016; Rylaarsdam & Guemez-Gamboa, 2019; Pavlinek et al., 2024). Similarly, the observed lower penetrance in the maternal cell line may exemplify the female protective theory, suggesting that females might require a more substantial accumulation of etiological factors to manifest autistic traits (LaSalle, 2013; Zhang et al., 2020; Cameli et al., 2021; Wigdor et al., 2022). These findings highlight the critical need to explore the complex interactions between genetic and sex-linked variants and *NRXN1* intronic deletions to understand how *NRXN1* influences molecular and functional changes, especially in individuals with similar deletions who display significantly different phenotypes.

In conclusion, deletions within intron 5 of *NRXN1* have a profound impact on the expression of *NRXN1* isoforms and are linked to disruptions in critical molecular mechanisms governing synaptic functionality and dendritogenesis. These differences manifests with a diversity in the expression of molecular and cellular phenotypes, suggesting that individual genetic makeup plays a crucial role in modulating the neurobiological outcomes of these deletions.

## Acknowledgements

This work was supported by a research grant from the Autism Research Trust (DPS, DA, SB-C) and University of Pennsylvania Autism Spectrum Program of Excellence (DPS, ZZ, MB). This work was also supported by grants from UK Medical Research Council, Grant No. MR/L021064/1, MR/X004112/1 and MR/Y012968/1to DPS. DPS acknowledges support from the UK Medical Research Council Centre for Neurodevelopmental Disorders (Grant No. MR/N026063/1); DPS is also a recipient of an Independent Researcher Award from the Brain and Behavior Foundation (formally National Alliance for Research on Schizophrenia and Depression) (Grant No. 25957). This work was supported by grants from The Simons Foundation Autism Research Initiative (DPS, DA, SB-C), the Autism Centre for Excellence (DA, SB-C). This study was supported by grants from the European Autism Interventions (EU-AIMS) and the Innovative Medicines Initiative Joint Undertaking under Grant No. 115300, resources of which are composed of financial contribution from the European Union’s Seventh Framework Programme (FP7/2007-2013) and the European Federation of Pharmaceutical Industries and Associations companies’ in kind contribution (DPS, SB-C); and Stem cells for Biological Assays of Novel drugs and prediCtive toxiCology (StemBANCC): support from the Innovative Medicines Initiative joint undertaking under Grant No. 115439-2, whose resources are composed of financial contribution from the European Union (FP7/2007-2013) and European Federation of Pharmaceutical Industries and Associations companies’ in-kind contribution (DPS). The authors acknowledge use of King’s Computational Research, Engineering and Technology Environment (CREATE) and are grateful to Dr George Chennell of the Wohl Cellular Imaging Centre at King’s College London for technical support with imaging.

## Author contributions (CRediT)

Lucia Dutan: Investigation, Conceptualization, Formal Analysis, Writing – original draft, Writing review & editing;

Nicholas J. F. Gatford: Investigation, Conceptualization, Formal Analysis, Writing – original draft, Writing review & editing;

Roland Nagy: Investigation, Methodology;

Thaise Carneiro: Investigation, Formal Analysis;

Victoria Higgs: Investigation, Formal Analysis;

APEX Consortia: Methodology;

Aicha Massrali: Methodology;

Arkoprovo Paul: Methodology;

Maria Fasolino: Methodology;

Zhaolan Zhou: Methodology;

Frances A Flinter: Resources;

Dwaipayan Adhya: Funding acquisition, Methodology, Writing review & editing;

Maja Bucan: Funding acquisition, Methodology, Writing review & editing;

Simon Baron-Cohen: Funding acquisition, Methodology, Writing review & editing;

Deepak P. Srivastava: Funding acquisition, Project administration, Supervision, Methodology, Writing – original draft, Writing review & editing.

## List of abbreviations

hiPSC: Human Induced Pluripotent Stem Cell
NRXN1: Neurexin 1
NRXN1α: Neurexin 1 alpha isoform
NRXN1β: Neurexin 1 beta isoform
2i: Dual SMAD inhibition
RNAseq: RNA sequencing
DEGs: Differentially expressed genes
GO: Gene Ontology

