## Supplementary Figure for "Impact of intragenic *NRXN1* deletions on early cortical development"

### Supplemental information

### Supplementary Figures

**A**

| Oligo name | Target | Oligo sequence (5' to 3') |
|---|---|---|
| NRXN1α F | Exon 7- Exon 8 | TGTGGACTTCCAGAGAGACG |
| NRXN1α R | Exon 8 | GGAAGACAAGGCCAGCTTTA |
| NRXN1β F | Intron 17 (NRXN1α) | ACCACATCCACCATTTCCAT |
| NRXN1β R | Exon 18 (NRXN1α) | ATTTGTCCACCACCTTTGCT |

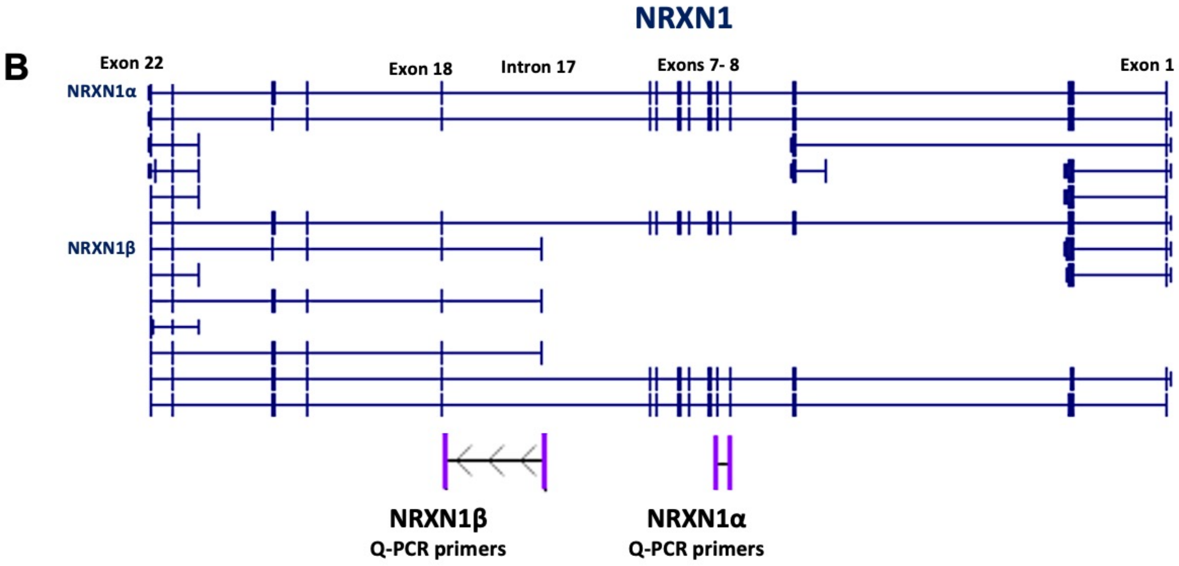


**Supplementary Figure 1:** Quantitative PCR Primers for NRXN1α and NRXN1β. (A) Primer Design and Specificity: This panel details the target locations and sequences of the primers used for NRXN1α and NRXN1β. The forward primer for NRXN1α spans exons 7 to 8, while the reverse primer is specific to exon 8. In contrast, the forward primer for NRXN1β targets intron 17, and the reverse primer targets exon 18 of the NRXN1α isoform. (B) Isoform Mapping: Displays a schematic map of the NRXN1 isoforms with genomic coordinates referenced from the UCSC Genome Browser (GRCh38/hg38), illustrating the specific regions targeted by the primers used in our study.

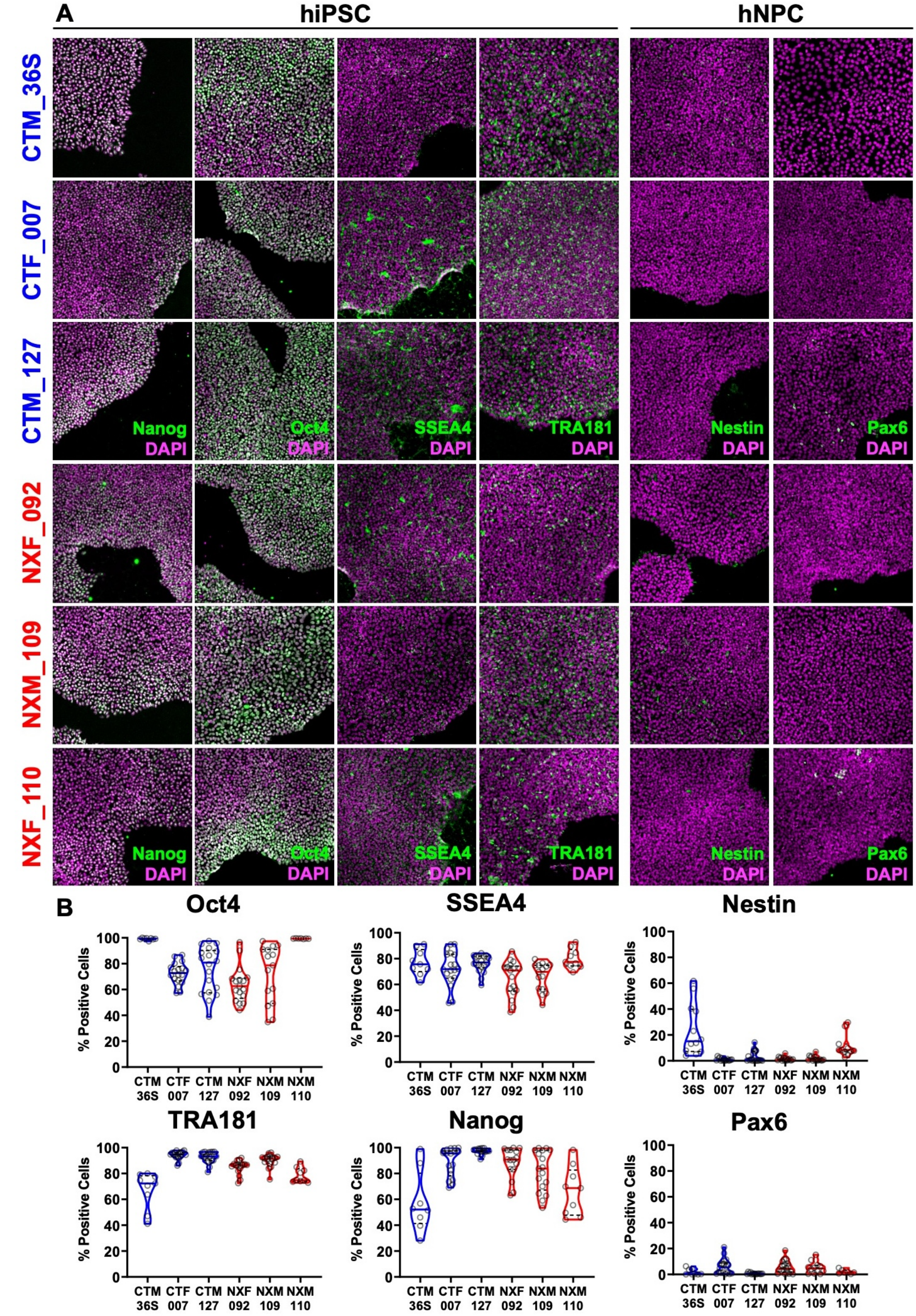
A
hiPSC
hNPC
CTM_36S
CTF_007
CTM_127
NXF_092
NXM_109
NXF_110
Nanog DAPI
Oct4 DAPI
SSEA4 DAPI
TRA181 DAPI
Nestin DAPI
Pax6 DAPI
B
Oct4
SSEA4
Nestin
TRA181
Nanog
Pax6
% Positive Cells
CTM 36S
CTF 007
CTM 127
NXF 092
NXM 109
NXM 110

**Supplementary Figure 2:** Assessment of Pluripotency and Neural Progenitor Cell Markers in hiPSCs. **(A)** Representative immunofluorescence images of human induced pluripotent stem cells (hiPSCs) at day 0, stained for pluripotency markers NANOG, OCT4, SSEA4, and TRA-1-81, alongside neural progenitor cell (NPC) markers NESTIN and PAX6. Images showcase both control lines (36S_CTR, 007_CTF, 127_CTM) and NRXN1 deletion lines (092_NXF, 109_NXM, 110_NXF), demonstrating the maintenance of pluripotency without notable spontaneous differentiation. **(B)** Violin plots illustrating the distribution of cells positive for pluripotency and NPC markers across control and NRXN1 deletion lines. Each plot displays median values (solid blue/red lines for control and deletion lines, respectively) and interquartile ranges (dotted black lines).

All scale bars represent 100 µm.

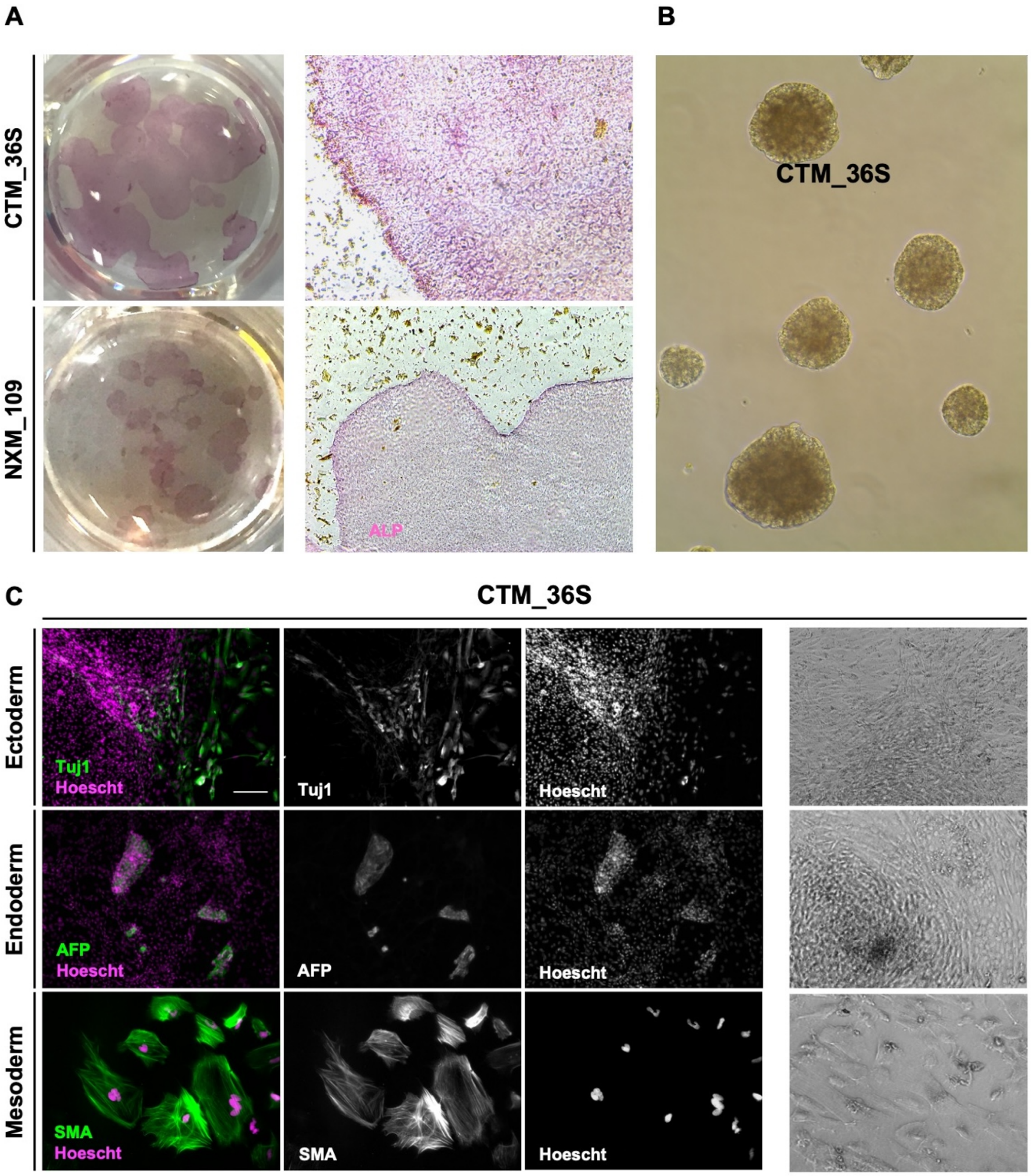


**Supplementary Figure 3:** Validation of Pluripotency in Control and NRXN1-Deletion Cell Lines. **(A)** Representative images of hiPSC colonies from the control line CTR_M3 and the NRXN1 deletion line 109_NXM stained for alkaline phosphatase activity, a marker of pluripotency, highlighting the enzymatic activity on the cell surface. **(B)** Brightfield microscopy image of the control line CTR_M3 hiPSCs after 48 hours in suspension culture, showcasing the capability of forming embryoid bodies, a characteristic feature of pluripotent stem cells. **(C)**

Combined fluorescence and phase-contrast imaging of CTR_M3 hiPSCs: **Left:** Fluorescence images showing positive staining for markers representative of the three germ layers: Tuj1 (ectoderm), AFP (endoderm), and SMA (mesoderm), confirming the pluripotency of the cells. Scale bar = 200 µm. **Right:** Phase-contrast images illustrating a variety of cellular morphologies corresponding to different germ layers, including fibrous ectoderm-like cells, small, rounded endoderm-like cells, and large flat mesoderm-like cells.

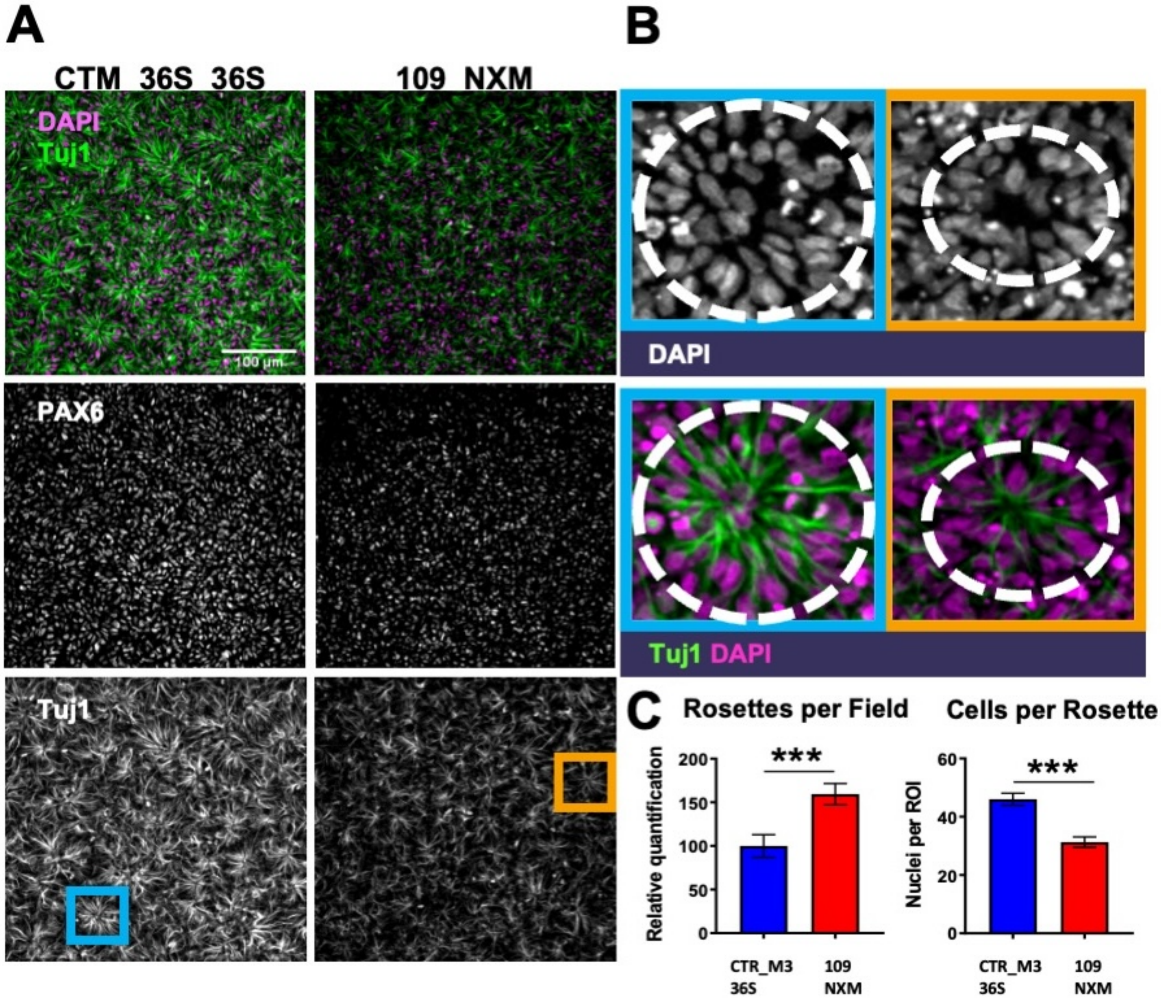


**Supplementary Figure 4:** Analysis of Neural Rosette Formation at Day 8. (A) This panel demonstrates the expression of the nuclear marker DAPI (magenta) and the neuronal processes marker Tuj1 βIII-tubulin (green), used to assess rosette formation in the control cell line CTR_M3_36S (left) and NRXN1 deletion line 109_NXM (right). (B) Representative images display the radial arrangements in both54r45e cell lines, highlighting that rosettes in the 109_NXM cell line are smaller compared to the control. Scale bars= 100 μm (10X objective) (C) Relative quantification of neural rosettes per field (arbitrary units) was determined by quantification of TUJ1 staining per field, ***$p$=0.0001 and quantification of cells contributing to rosettes was assessed by manual regions of interest (ROI) allocation using TUJ1 staining and quantification of nuclei per ROI ****$p<0.0001$.
